# Monitoring fatty acid trafficking during *Drosophila melanogaster* oogenesis reveals a role for the triglyceride synthase DGAT1 in protecting mitochondrial integrity

**DOI:** 10.64898/2026.08.10.743872

**Authors:** Roger P. White, Marcus D. Kilwein, Michael A. Welte

## Abstract

Successful oogenesis requires the precise coordination of nutrient uptake, storage, and utilization to meet the high metabolic demands of egg production. In mammals, fatty acid (FA) metabolism has emerged as a key driver of oocyte maturation; however, the mechanisms by which follicles regulate FA trafficking and utilization remain poorly understood across all systems. To address these issues, we leverage the genetic tractability of *Drosophila melanogaster* oogenesis. We found that nurse cell mitochondria are metabolically active and catabolize FA in a stage-dependent manner, with fatty acid oxidation (FAO) peaking during mid-oogenesis. By exposing explanted follicles to fluorescently labeled FAs, we monitored FA trafficking and found massive enrichment in lipid droplets. Mutants for the triglyceride lipase ATGL exhibited a reduction in both mitochondrial membrane potential and FAO, suggesting that mitochondria utilize FA from triglycerides stored in LDs. To determine the significance of this transient FA storage in LDs, we prevented the formation of nurse cell LDs with mutations in the triglyceride synthase DGAT1. The *DGAT1* mutant follicles display excess accumulation of FAs in mitochondria, mitochondrial stress, and developmental arrest. We find that this mitochondrial dysfunction and follicle arrest are consequences of FA toxicity to mitochondria: limiting FA influx into follicles or FA import into mitochondria alleviates these defects. Our findings demonstrate that LD-derived FAs are actively mobilized to fuel the energy demands of oogenesis while LDs buffer against lipotoxicity, revealing a critical balance between FA storage and oxidation. These findings highlight LDs as central hubs regulating energy homeostasis and developmental progression in the follicle.

**Author Summary:** Oogenesis places extraordinary metabolic demands on the follicle, yet the energy source for follicle development is not well understood. In fruit flies, follicles take in large amounts of lipids from the hemolymph, the insect blood, and accumulate massive fat stores in the form of lipid droplets. Whether these stores are reserved for the embryo or already power oogenesis was unclear. Using mutants and fluorescent probes for metabolic activity, we found that some fatty acids are released from the lipid droplets and power energy production in mitochondria; this energy source is important for successful oogenesis. We then fed flies fluorescently labeled fatty acids and determined how these fatty acids travel when lipid droplet formation can occur versus when it is abolished. In the former case, fatty acids accumulate in lipid droplets; in the latter, they flood into mitochondria, causing mitochondrial dysfunction, reduced ATP levels, and follicle death. We can correct all these defects by limiting lipid influx specifically into mitochondria. Our findings reveal an important role for lipid droplets during oogenesis. They act as a metabolic buffer, supplying sufficient amounts of fatty acids to mitochondria for energy production while shielding the mitochondria from toxic lipid levels.

## Introduction

Successful reproduction depends on the ability to produce an egg capable of fertilization and subsequent embryonic development [1–3]. Developing oocytes undergo massive growth (e.g., 500-fold increase in volume for human oocytes) and dramatically restructure their metabolism, accumulating an abundance of organelles and nutrients [4]. Even though all these changes are presumably extremely energy demanding, it remains largely unclear what energy sources promote oocyte maturation [1, 5].

Much of what is known about the metabolic needs of oogenesis comes from studies on mammalian follicles. These follicles contain an immature oocyte surrounded by somatic follicular cells that protect and nourish the oocyte [1, 5]. In vitro, oocytes devoid of these follicular support cells can nevertheless mature if the culture medium provides the required nutrients [1, 5]. Such studies have revealed a critical role for fatty acids (FAs) as well as for their catabolic breakdown via FA oxidation (FAO), a process that occurs in mitochondria. For example, in mouse, bovine, and porcine oocytes, pharmacological inhibition of FAO impairs oocyte maturation [1, 2, 6].

These studies also revealed that elevated concentrations of saturated FAs are detrimental to oocyte developmental competence, while monounsaturated FAs have protective or beneficial effects on oocyte quality [2, 3, 6]. These observations suggest that during oogenesis, both the quantity and type of FAs must be tightly regulated [1, 5]. However, while these experiments demonstrate the importance of FAs and FAO for follicle development, it remains unknown how FAs are trafficked into follicles, how they are utilized, and how dysregulated FA metabolism influences the success of oogenesis.

To start addressing these questions, we are taking advantage of the well-characterized oogenesis of *Drosophila melanogaster*. The *Drosophila* follicle is composed of an outer layer of somatic epithelial cells known as follicle cells, which enclose a cyst of sixteen interconnected germline cells: one oocyte and fifteen nurse cells (NCs) [7]. The roughly week-long development of the follicle is divided into 14 stages, S1 to S14 [8]. From S9 to S14, the oocyte undergoes massive growth, increasing its volume ∼100 fold in just 3 days, via two mechanisms. First, a large portion of the eventual ooplasm is generated in the polyploid NCs and then transferred to the oocyte [9, 10]; this material includes RNAs and proteins as well as organelles, such as the ER, mitochondria, and LDs, the cellular sites for fat storage [11]. Second, the follicle imports materials from the hemolymph, the insect blood, including massive amounts of both proteins and lipids: yolk proteins originate in the fat body and are taken up by oocytes via endocytosis [11, 12]; diacylglycerol-rich lipophorin particles transport lipids from the gut and fat body to the NCs to support LD formation [13].

During germarial and early pre-vitellogenic stages (through S4), NC metabolism relies on a balance of oxidative and glycolytic metabolism [14]. Oxidative metabolism continues to be a source of energy until late oogenesis (S13-14) [14–16] when mitochondria become respiratory quiescent [15]. After S8, it is unlikely that oxidative phosphorylation is mainly fueled by glucose, as glucose is preferentially funneled into the pentose phosphate pathway to drive nucleotide synthesis to sustain high rates of DNA replication [17]. Indeed, ablation of key enzymes in this pathway results in reduced egg production while interfering with glycolysis does not [17]. Thus, the energy needs of mid to late oogenesis must be met in some other way. Whether this energy demand is met by FAO as in mammals has not been critically evaluated.

In principle, LDs could provide substrates for FOA as they are well known to play such a role in other tissues and are particularly abundant in follicles. For instance, in mouse skeletal muscle, triglycerides in LDs are hydrolyzed by Adipose Triglyceride Lipase (ATGL) to release free FAs, which are subsequently oxidized via FAO in mitochondria to generate ATP [18, 19]. During fly oogenesis, early-stage follicles contain a smattering of LDs, but during S9, LDs accumulate rapidly, and by S10B, NCs are densely packed with hundreds of thousands of LDs [9, 11, 20]; most of these LDs are transferred to the oocyte during S11. However, whether these LDs are employed for energy production remains an open question. On the one hand, the LDs generated during oogenesis may need to be preserved since they are a critical reserve to power the energy needs of embryonic development [21]. On the other hand, ATGL is active to liberate arachidonic acid from NC LDs, thereby promoting prostaglandin synthesis [10, 20]. It is thus conceivable that LD-derived FAs also serve as substrates for mitochondrial FAO and support oogenesis.

In this study, we find that in mid-oogenesis, NC mitochondria indeed rely on FAO to bolster their membrane potential; the fatty acids used as fuel are, in part, released from LDs by ATGL. To visualize how FAs traffic through follicles in real time, we supplied follicle explants with fluorescently labeled FAs. In the wild type, these FAs predominantly accumulate in LDs; in mutants lacking the triglyceride synthase DGAT1, no LDs are produced, and these FAs massively localize to mitochondria. *DGAT1* mutant follicles display high levels of mitochondrial ROS and die by apoptosis in S9. ROS levels and follicle death are markedly reduced when FA import is decreased, either into the follicle as a whole or into mitochondria specifically. We conclude that a critical role of NC LDs is to protect mitochondria from FA overload, thus ensuring the production of healthy oocytes.

## Results

### *Drosophila* Follicles Exhibit Fatty Acid Oxidation

To investigate whether FAO occurs at all in follicles, we employed the compound FAOBlue, a FA conjugated with the dye coumarin; in this form, the dye is non-fluorescent. In mitochondria, FAOBlue is metabolized like a FA and broken down by β-oxidation, releasing free, now fluorescent, coumarin into the cytoplasm [22]. After dissecting ovaries out of females, we incubated follicles of various stages ex vivo with FAOBlue (Fig 1A-A’); follicles incubated without the compound served as a control (Fig 1B-B’). We then quantified the FAOBlue fluorescence for various follicle stages. We detected by far the strongest signal in S9 and S10 follicles; in previtellogenic stages (S1-S8), the signal was low to undetectable (Fig 1A-A’, A’’). In S9 and S10 follicles, it was mostly the cytoplasm of NCs that was FAOBlue positive, with little signal in oocytes or follicle cells (Fig 1A-B; the oocyte signal visible in Fig 1A is due to yolk vesicle autofluorescence and is already apparent without FAOBlue). This pattern is intriguing as the NCs of S9 and S10 highly upregulate the expression of lipophorin receptors [23] and thus likely have tremendous FA influx from the hemolymph. High FA intake is also suggested by the fact that in these stages NCs accumulate large amounts of LDs [20].

**Fig 1.**
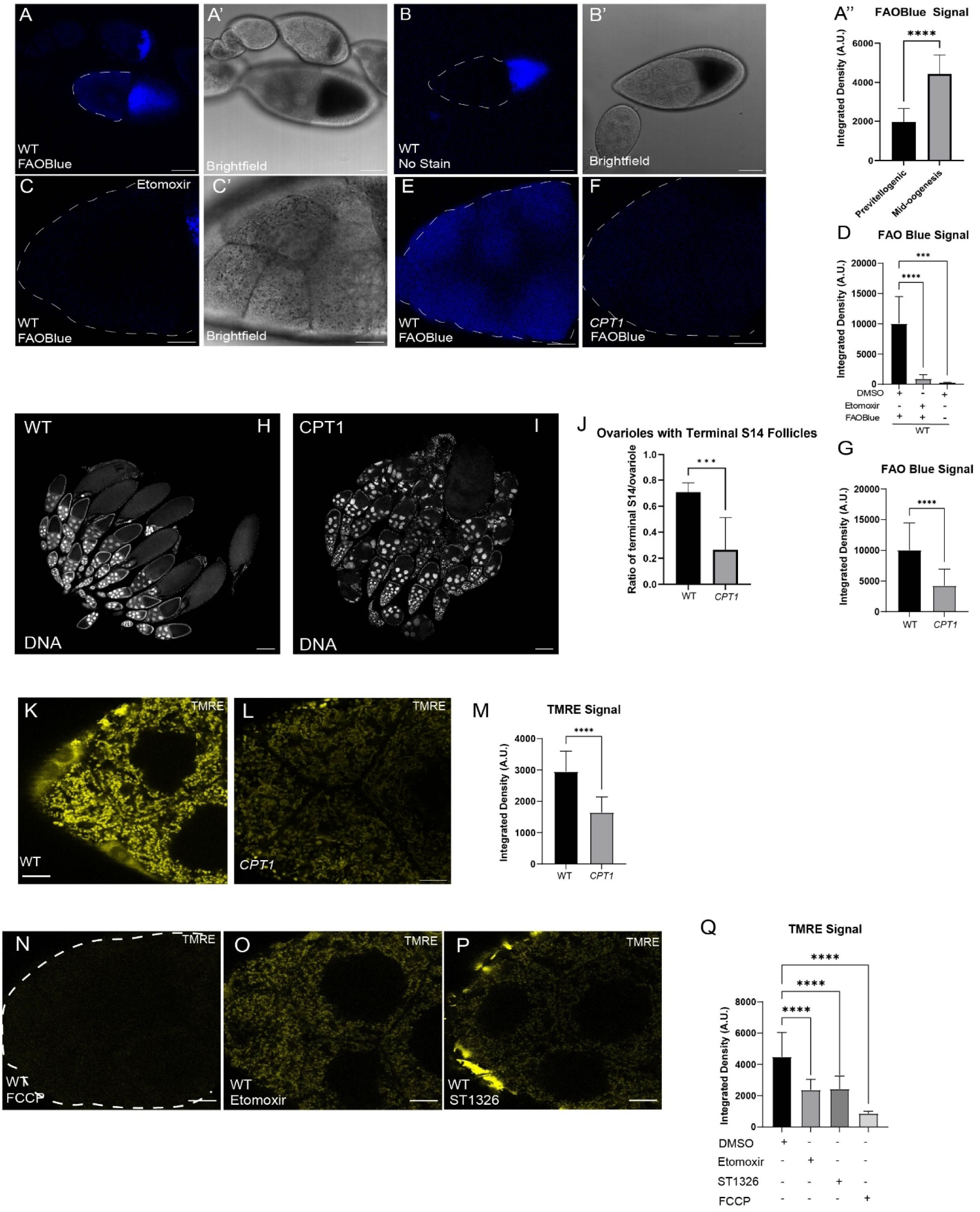
*Drosophila* Follicles Undergo Fatty Acid Oxidation. **A-A’)** FAOBlue staining (A) and brightfield image (A’) of a section of a wild-type ovariole. A’’) Quantitation of FAOBlue signal of previtellogenic follicles (S1-S8) and mid-oogenesis follicles (S9). Relative fluorescence was quantified using a defined ROI to record the integrated density. B-B’) Explanted wild-type follicles in the absence of FAOBlue stain imaged for FAOBlue signal (B) or by brightfield microscopy (B’). B was imaged using the same imaging parameters as for A; the signal from oocytes is autofluorescence from yolk vesicles. C-C’) Nurse cells of explanted wild-type S9 follicle treated with 50μM etomoxir prior to incubation with FAOBlue; FAOBlue signal (C) and brightfield image (C’). D) Quantitation of FAOBlue signal from wild-type follicles with various drug treatments. E-F) FAOBlue staining of explanted S9 wild-type (E) and *CPT1* mutant (F) follicles. G) Quantitation of FAOBlue signal in wild-type and *CPT1* mutant S9 follicles. H) Tiled images of whole wild-type and I) *CPT1* mutant ovaries stained with Hoechst (DNA). J) Quantitation of ovarioles with terminal S14 follicles. K) TMRE staining of wild-type and L) *CPT1* mutant S9 follicles. M) Quantitation of TMRE integrated density normalized to MitoTracker green signal for wild-type and *CPT1* mutant S9 follicles. N) Wild-type S9 follicles treated with etomoxir, O) ST1326, or P) FCCP, respectively, and then stained with TMRE. Q) Quantitation of TMRE integrated density normalized to MitoTracker green signal for various drug treatments. Scale bars in A = 50μm. Images were acquired on Leica Stellaris (20x) for panels H-I only Scale bar in B-E = 20μm. Scale bar in G-H = 100 and 150μm, respectively. Scale bar in J-K and M-O = 10μm. \*\*\**p<0.001, ****p<0.0001*

If the FAOBlue signal indeed faithfully reads out FAO activity, it should be sensitive to the amount of FAs reaching mitochondria. We therefore exposed follicles to etomoxir to inhibit the enzyme carnitine palmitoyltransferase 1 (CPT1/Withered) [24], responsible for the import of long-chain FAs into mitochondria [25]. With drug treatment, FAOBlue signal was almost abolished, similar to follicles not treated with the compound (Fig 1C-D). We conclude that FAO is indeed active during *Drosophila* oogenesis, in a stage and cell-type-specific manner.

### *CPT1* Mutants Exhibit Oogenesis Defects and Reduced Mitochondrial Membrane Potential

To determine if FAO makes important contributions to oogenesis, we analyzed a null allele in *CPT1* [25]. Follicles from *CPT1* mutant females had reduced FAOBlue signal, consistent with acute inhibition of CPT1 in wild-type flies (Fig 1E-G). Compared to wild-type ovaries, those from *CPT1* mutant females had very few late-stage follicles, and many S9 and S10 follicles showed developmental abnormalities (Fig 1H-J), including condensation of NC nuclei, failure of follicle cells to undergo elongation, reduced oocyte size in S10, and NC blebbing. Together, these observations suggest that lack of FAO compromises oogenesis.

We therefore probed the bioenergetics of NC mitochondria using the dye Tetramethylrhodamine ethyl ester (TMRE). TMRE is a cell-permeant, cationic fluorescent dye that is readily sequestered by polarized mitochondria and thus can be used to assess the mitochondrial membrane potential (MMP) [26, 27]. We explanted S9 follicles into media containing TMRE and imaged them live by fluorescence microscopy. TMRE uptake into mitochondria occurred rapidly for both NCs and follicle cells, revealing membrane potential and morphology of the mitochondria. Signal was apparent with just 15 minutes of incubation and remained largely unchanged even after an hour (S1A-B Fig). This stability allowed us to assess the effects of various pharmacological treatments on signal strength in later experiments. As a control, we abolished the MMP using the uncoupler Carbonyl cyanide 4-(trifluoromethoxy) phenylhydrazone (FCCP) [26, 27]. FCCP treatment completely ablated the TMRE signal in all follicle stages (Fig 1N, Q).

Using the same assay, we examined the mitochondrial membrane potential (MMP) in *CPT1* mutant follicles. There were no striking differences from wild type in early follicle stages, and in S10, follicle cells and oocytes displayed a very similar signal to the wild type. However, the signal in NCs was significantly reduced (Fig 1K-M). Such a drop in TMRE signal could be explained if NC mitochondria have a lower membrane potential or if there were changes in mitochondrial mass. We therefore co-stained with the dye MitoTracker Green, which accumulates in mitochondria independent of their membrane potential, and computed the ratio of the two dyes [26, 27]. This analysis revealed that changes in mitochondrial mass were not the reason for TMRE differences but rather reflect a reduction in MMP (S1C-D Fig). Consistent with this interpretation, we observed a similar drop in TMRE signal in the wild type with acute inhibition of CPT1, using two different inhibitors, etomoxir and ST1236 (Fig 1O-Q). Treatment with the vehicle alone did not exhibit any changes in signal (S1A-B Fig). In summary, we conclude that the lack of CPT1 activity results in reduced MMP, specifically in NCs of stages with high FAO. It also compromises the production of developmentally competent follicles.

### Monitoring Fatty Acid Flux into Follicles

The FAs used in NC mitochondria might be produced de novo in the NCs themselves or might originate in other cell types or tissues. To monitor how exogenous FAs are incorporated into follicles, we fed flies for 20 hours with yeast paste supplemented with a fluorescently labeled fatty acid, namely palmitic acid conjugated with the fluorescent dye BODIPY (C16:0-BODIPY). As the dye is attached to the hydrocarbon tail distal to the carboxyl group, it can be incorporated into cellular lipids just like the native FA [28]. Similar fluorescent FAs (FLFAs) have been employed in cultured cells to monitor FA trafficking [28]. After 20hr feeding, we dissected the ovaries, fixed them, and examined the distribution of FLFAs by confocal imaging. We detected fluorescent signal in distinct puncta in the NCs and follicle cells (S2A-A’’ Fig). This assay can be used to observe the accumulation of different types of FAs into follicles; for example, we previously published similar data using fluorescent arachidonic acid, a polyunsaturated FA [20].

Presumably, the FLFA taken up through the gut is trafficked, directly or indirectly, via lipophorin particles to the ovary. Previous studies have established a general framework for how FAs travel to follicles. Lipophorin particles from the hemolymph, rich in diacylglycerides (DAG), dock to lipophorin receptors LpR1 and LpR2 at the surface of NCs. These receptors, as well as the lipophorin co-factor LTP, are required for efficient LD accumulation in NCs [13, 23], consistent with the idea that this pathway is crucial for transferring FAs to NCs.

To monitor FA influx in real time, we incubated explanted follicles in media with DAG containing a FLFA (C16:0-BODIPY/C11:0) and found that fluorescence was readily taken up by the follicle (S2B-B’ Fig). These conditions presumably mimic lipid delivery via the DAG-rich lipophorin particles. As it was proposed that the breakdown of DAG occurs extracellularly [13, 23], and it is therefore FAs that are taken up by the NCs, we tested whether we could even circumvent the lipolysis step and provide FAs directly to the follicles. Indeed, incubating follicles with C16:0-BODIPY resulted in the accumulation of fluorescent puncta in the NC cytoplasm and follicle cells (Fig 2A-C). Fluorescent puncta were detectable within 15 minutes in the NC cytoplasm; their intensity initially increased and then seemed to level off; over time, more puncta appeared, until they filled the entire NC cytoplasm. Closer inspection and simultaneous detection of LDs revealed that these puncta are LDs and that label accumulation was uneven, with some LDs accumulating label to much higher levels than others (S1 Video). We presume that some FLFAs are incorporated into already existing LDs, while LDs newly generated during the incubation period reach much higher levels of fluorescence. We tested to see if incorporation of FLFA into LDs is an active process by fixing follicles prior to incubation and failed to detect substantial accumulation (S2C-D’’’ Fig); thus, FLFA accumulation is not due to simple passive partitioning to the LD neutral lipid core. We observed similar gradual puncta formation when using other FLFA and sterols, such as BODIPY C4, C9, arachidonic acid (C20:4-NBD), and cholesterol (25-NBD Cholesterol) (S2E-G Fig). We conclude that we can monitor the flux of FAs and sterols into follicles via live imaging and that these lipid species predominantly accumulate in LDs.

**Fig 2.**
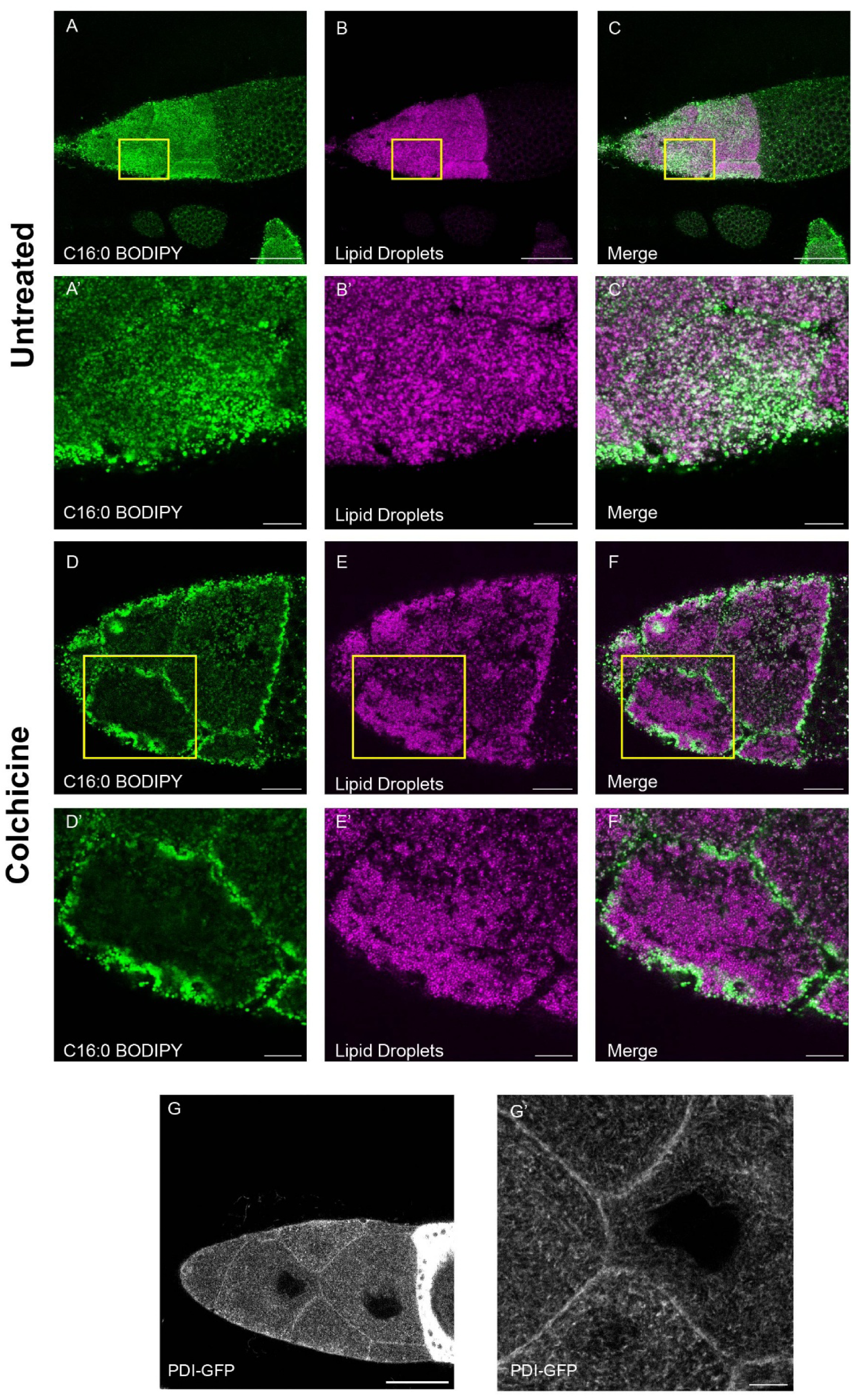
NC LDs Incorporate FLFAs and Are Born Near the ER at the NC Cortex. A-C’) Untreated (No colchicine) wild-type S10 follicle after 20-minute incubation with 5μM C16:0-BODIPY and then fixed with 4% paraformaldehyde. BODIPY signal (A), LD staining (LipidSpot) (B), and merge of BODIPY and LD channel (C). A’, B’, and C’ are magnified regions of the same images. D-F’) S10 Follicle from wild-type flies that had been fed 80μg/mL colchicine yeast paste for 6h. After dissection, follicles were incubated with C16:0-BODIPY for 20 minutes and fixed with 4% paraformaldehyde. BODIPY signal (D), LD staining (LipidSpot) (E), and merge of BODIPY and LD channel (F). D’, EB’, and FC’ are magnified regions of the same images. G) S10 Follicle expressing PDI-GFP, fixed with 4% paraformaldehyde. G’) Magnified region from G. Scale Bar A-C = 50 μm A’-C’ = 10 μm. Scale Bar for D-F = 20 μm scale Bar D’-F’ = 10 μm Scale Bar G = 50 μm G’ = 10 μm

### NC Lipid Droplets are Highly Motile and are Born near the cortical ER

We next monitored the time course of FLFA accumulation in LDs by taking movies of S9 follicles exposed to C12:0-BODIPY in the media. This live imaging revealed that FLFA puncta move rapidly within the NC cytoplasm (S1 Video). Individual puncta moved linearly for many seconds, but there was no obvious directionality of the population as a whole. The net results appeared to be a rapid mixing of the LDs. To determine if this motility is a general feature of all LDs, we took movies by confocal reflection microscopy to visualize LDs and found the same overall pattern (S2 Video). Directed motion of LDs had previously been observed in S6 NCs [29], in S12 oocytes [30], and early embryos [21, 31]. While the exact mechanisms of motility differ between these stages, they all rely on cytoskeletal elements, including cytoplasmic streaming powered by actin or microtubule-based motors, microtubule gliding, or bidirectional transport of LDs along microtubules. Microtubules also make an important contribution to LD motility in S9/10 NCs, as treatment of follicles with the microtubule depolymerizing drug colchicine largely abolished motion (S3 Video). LDs immobilized after colchicine treatment tended to cluster together (Fig 2D-F’).

When we incubated follicles with C12:0-BODIPY after colchicine treatment, the distribution of fluorescent puncta was highly polarized. Almost all the puncta were closely associated with the NC cortex rather than dispersed through the NC cytoplasm (Fig 2D-F). These data suggest that new LDs are typically generated at the NC cortex. Presumably, FAs delivered at the NC plasma membrane are rapidly packaged into LDs, rather than reaching other organelles where a massive, unregulated FA influx could cause lipotoxicity. LD biogenesis occurs at the endoplasmic reticulum (ER), where newly synthesized neutral lipids first accumulate in between ER leaflets and then come together to form a nascent LD [32]. Therefore, if LDs are indeed generated at the NC cortex, the cortex should be rich in ER. Using the ER marker PDI-GFP, we indeed found that the ER is present along the NC cortex in mid-oogenesis follicles (S9-10) (Fig 2G-G’). These findings are consistent with other reports that, using the ER marker Calnexin, found high accumulation of ER at the NC periphery in S10B [20, 33].

### Normal Mitochondrial Membrane Potential Requires the Triglyceride Lipase ATGL

Our analysis of *CPT*1 mutants strongly argues that NC mitochondria rely heavily on FAO, but we have not been able to detect any significant accumulation of FLFAs in mitochondria, presumably because they are turned over rapidly there (Fig 3A-A’). This pattern does not allow us to distinguish whether FAs newly arrived in follicles directly travel to mitochondria or whether they are transiently stored in LDs before being delivered to mitochondria. It has indeed been suggested that in many cells, LDs serve as central hubs for FA trafficking and gather FAs from diverse locations before funneling them into mitochondria [11, 28].

**Fig 3.**
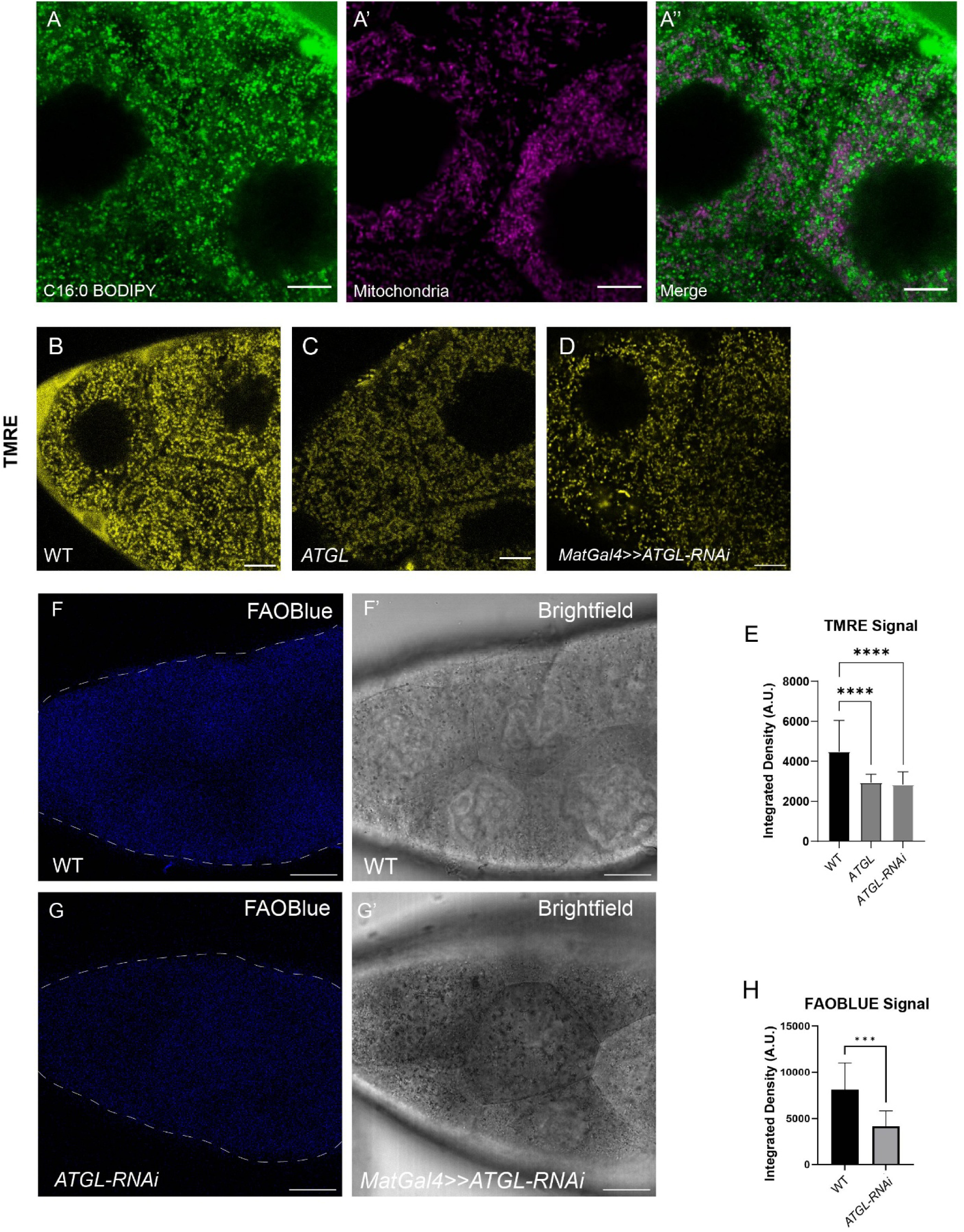
Normal Mitochondrial Membrane Potential Requires the Triglyceride Lipase ATGL. **A-A”)** Live image of wild-type S9 follicle supplemented with 5 μM C16:0-BODIPY (A) and stained with MitoView (A’); A’’ is the merge of the two channels. Still was captured 30 minutes after introduction of C16:0-BODIPY B-D) Live images of wild-type (B), *ATGL^1^* (C), and *ATGL-RNAi* (D) S9 follicles after 15-minute incubation with TMRE (25nM). The maternal tubulin Gal4 driver was used to drive the expression of ATGL-RNAi in NCs. E) Quantitation of the integrated density of the TMRE signal for S9 follicles of the indicated genotypes. F-F’) Live image of FAOBlue staining (F) and the corresponding brightfield image (F’) of a wild-type S9 follicle. G-G’) Live image of FAOBlue staining (G) and the corresponding brightfield image (G’) of an *ATGL-RNAi* S9 follicle. H) Quantitation of FAOBlue staining S9 for follicles of the indicated genotypes. Scale Bar A-A’’ = 10μm. Scale Bar B-D = 10 μm. Scale Bar F-G’ = 20 μm.

If FAs are transiently stored in triglycerides first, their release into mitochondria would require lipolytic breakdown. In many cells, the major triglyceride lipase is Adipose Triglyceride Lipase (ATGL, also known as Brummer in flies) [34]. We have previously shown that ATGL is expressed in NCs and localizes to LDs [20]. It must also be active there, because it is required for the release of AA from triglycerides and its use as a substrate for prostaglandin biosynthesis [20].

We therefore examined the MMP in NCs of *ATGL* mutants. Compared to wild type, the MMP was significantly reduced (Fig 3B). This impairment is cell-autonomous as germ-line specific knockdown of ATGL resulted in a similar reduction of MMP (Fig 3C-E). Such knockdown also resulted in reduced FAOBlue signal (Fig 3F-H). These data suggest that ATGL has a role in regulating the release of FA from LDs and is a major regulator of the pool of FAs delivered into mitochondria for FAO.

### *DGAT1* Mutants Display Multiple Mitochondrial Defects

Our data argue that exogenous FAs are packaged into LDs and that mitochondrial FAO relies, at least in part, on FAs released from LDs. Why are FAs taking this circuitous route, being sequestered in LDs first before being delivered to mitochondria? To address this question, we utilized mutants for the triglyceride synthase, *DGAT1*. We confirmed previous findings that in strong loss-of-function alleles of *DGAT1,* NCs of S9 follicles contain very few LDs [9]. The LD content of follicle cells appears normal, suggesting that LD formation relies on a different neutral lipid synthesis pathway (Fig 4A-B). Most *DGAT1* mutant follicles die in S9 by apoptosis; because of this mid-oogenesis death, the gene is also known as *midway* [9, 35]. For the analysis below, we selected *DGAT1* follicles that did not yet show obvious signs of degeneration. In particular, we confirmed by brightfield imaging that NC nuclei did not appear condensed, that border cells were undergoing migration, and that follicle cells were elongating. Older S9 *DGAT1* follicles show more signs of defects and were generally avoided for experiments, unless specifically noted to record exacerbated damage.

**Fig 4.**
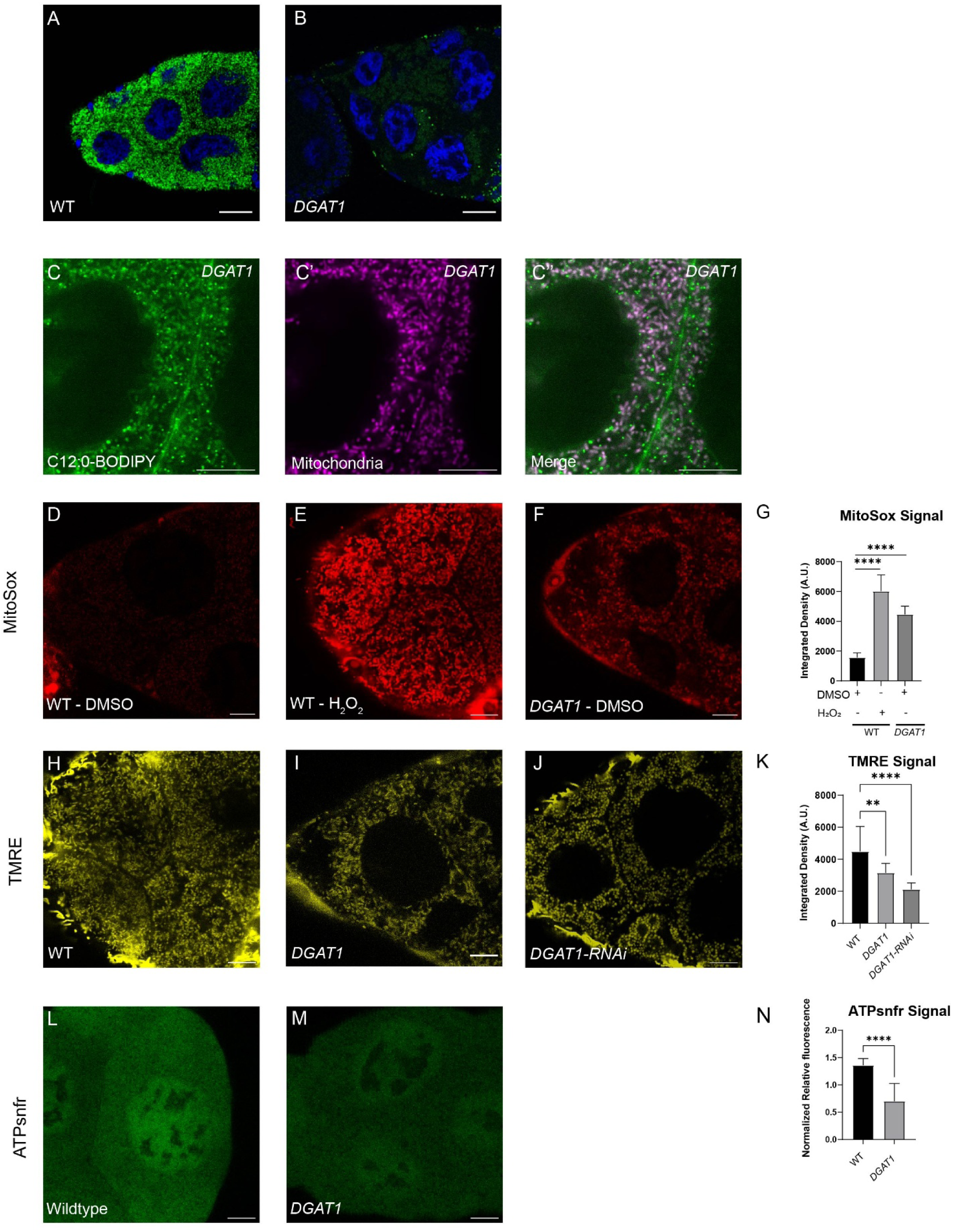
*DGAT1* Mutant Follicles Display Multiple Mitochondrial Defects. **A)** Fixed wild-type and B) *DGAT1* mutant S9/10 follicles stained with Nile Red and Hoechst. C-C”) Live images of *DGAT1* nurse cell supplemented with 5μM C16:0-BODIPY and stained with Mitoview. BODIPY signal (C), Mitoview signal (C’), merge of the two channels (C’’). Still was captured 30 minutes after the introduction of C16:0-BODIPY. D) Live image of wild-type, E) wild-type H_2_O_2_-treated, and F) *DGAT1* S9 follicles after 30-minute incubation with MitoSox. G) Quantitation of MitoSox signal for S9 follicles of the indicated genotypes. H-J) Live images after 15-minute incubation with TMRE (25nM) of a wild-type (H), *DGAT1* (I), or *DGAT1-RNAi* (J) S9 follicles. The maternal tubulin Gal4 driver was used to drive the expression of *DGAT1-RNAi* in NCs. K) Quantitation of TMRE signal for S9 follicles of the indicated genotypes. L) Images of ATP synthase sniffer signal in wild-type and M) *DGAT*1 follicles. N) Quantitation of normalized levels of ATP synthase sniffer fluorescence for S9 follicles of the indicated genotypes. Images were acquired on Leica Stellaris (63x) for panels C-C’’ only Scale Bar A-B = 20 μm. Scale Bar C-C’’ = 10 μm. Scale Bar D-F = 10 μm. Scale Bar H-J = 10 μm. Scale Bar L-M = 10 μm.

When we incubated *DGAT1* follicles ex vivo with the FLFA C12:0-BODIPY, fluorescent signal still appeared in distinct puncta as well as in a network of tubules that presumably represent the ER. Co-staining with the mitochondrial dye Mitoview revealed that these puncta are mitochondria (Fig 4C-C’’). Thus, when LDs cannot be formed, incoming FLFAs inappropriately accumulate in mitochondria, with some excess also in the ER.

While FAO is an important source of energy production, it also generates reactive oxygen species (ROS) and thus can result in oxidative stress. To test whether the increased mitochondrial FA levels in *DGAT1* mutants increase oxidative stress, we utilized the superoxide indicator MitoSOX; superoxide is a deleterious byproduct of mitochondrial respiration and is toxic in high concentrations. MitoSOX accumulates in mitochondria in an MMP-dependent manner and thus can be used to specifically detect ROS in mitochondria (mitoROS). As proof-of-principle, we treated wild-type follicles with hydrogen peroxide to induce mitoROS accumulation and compared them with follicles not exposed to the compound; treated follicles had over a three-fold increase in MitoSOX fluorescence (Fig 4D-E, G). Compared to untreated wild type follicles, untreated *DGAT1* mutant follicles exhibited a two-fold increase in MitoSOX fluorescence (Fig 4F-G), indicating that *DGAT1* mutant NC mitochondria are experiencing increases in mitoROS. In further support of this conclusion, incubating *DGAT1* mutant follicles with the ROS scavenger n-acetylcysteine amide (NACA) reduced MitoSOX levels (S3A-D Fig).

ROS has the potential to damage mitochondria and compromise their function. Indeed, TMRE signal of *DGAT1* follicles was significantly decreased, indicating compromised MMP (Fig 4H-K). Incubation with MitoTracker Green showed that mitochondrial mass was not different from wild type (S3E-F Fig). Therefore, the reduction in TMRE signal is not due to fewer mitochondria, but to a reduction in MMP. This defect in MMP is at least partially a result of excessive ROS production since treatment of *DGAT1* follicles with NACA restored MMP to near wild-type levels (S3G-J Fig). The reduction in MMP also raises a technical issue: the reagents for detecting ROS and MMP accumulate in mitochondria in a membrane potential-dependent manner; thus, the recorded MitoSOX signal in the *DGAT1* mutants underestimates ROS levels. We therefore computed the ratio between MitoSOX and TMRE signal, allowing us to estimate that relative to the wild type mitoROS levels are up ∼3-fold in the mutant (S3K Fig).

Do mitochondrial stress and MMP reduction impair mitochondrial function in ATP production? We took advantage of an in-vivo ATP sensor previously tested in cultured cells, ATP synthase sniffer (ATPsnfr) [36]. ATPsnfr consists of a microbial F_0_F_1_-ATP synthase epsilon subunit in which a circularly permuted GFP is inserted. ATP binding to the epsilon subunit results in a conformational change that increases GFP fluorescence. We generated a transgene encoding ATPsnfr under UASp control and drove it in the germline using maternal α-Tubulin Gal4. When we treated follicles with FCCP prior to imaging to stop ATP synthesis, we detected reduced GFP fluorescence compared to DMSO controls (S3L-N Fig). The biosensor is fused with RFP to allow for normalization. We quantified the ratio between GFP and RFP signals to normalize variations in expression levels of the construct. This experiment validates the use of ATPsnfr as an ATP sensor in NCs. Comparison of wild type and *DGAT1* mutants revealed that *DGAT1* mutants had lower relative ATPsnfr signal (Fig 4L-N, S3O-P Fig), consistent with the notion that mitochondrial damage compromises ATP production.

### Throttling FA Import into Mitochondria Ameliorates Many of the Mitochondrial Defects of *DGAT1* Mutants

Mitochondria in *DGAT1* mutants exhibit increased mitoROS and reduced MMP. We hypothesize that both defects are due to the excess accumulation of FAs in these mitochondria. To directly test this idea, we generated *DGAT1 CPT1* double mutants (Fig 5A-D). Lack of DGAT1 function will abolish NC LD formation, and lack of CPT1 should greatly reduce FA accumulation in mitochondria and thus ameliorate those phenotypes due to excess FAs in mitochondria.

**Fig 5.**
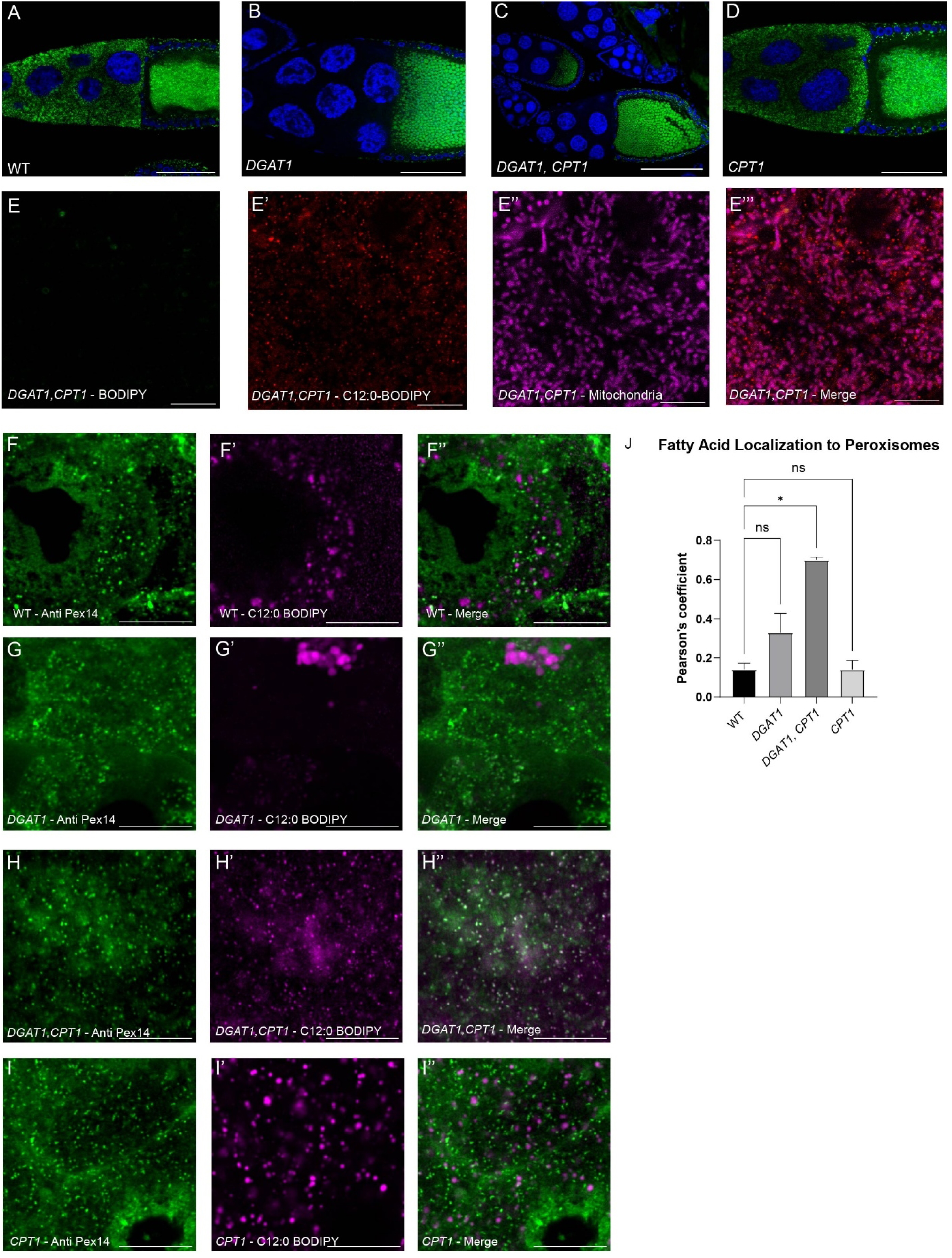
Throttling FA Import into Mitochondria Ameliorates many of the Mitochondrial Defects of *DGAT1* Mutants. A-D) Nile Red and Hoechst staining of fixed wild-type (A), *DGAT1* (B), *DGAT* CPT1 (C), and *CPT1* (D) mutant S9/10 follicles. E-E’’’) Live image of a *DGAT1 CPT1* S9 follicle that was stained with BODIPY (E), supplemented with 5μM C12:0-BODIPY (E’), and stained with Mitoview (E’’). E’’’) shows the merge of all channels. F-I”) S9 Follicles stained for PEX14 and supplemented with C12:0-BODIPY for wild type (F-F”), *DGAT1* (G-G’’), *DGAT1 CPT1* (H-H’’), or *CPT1* mutants. J) Quantitation of colocalization of C12:0-BODIPY and PEX14 immunostaining using Pearson’s coefficient for S9 follicles of the indicated genotypes. Colocalization was determined using the Imaris Coloc tool. Images were acquired on Leica Stellaris (63x) for panels E-E’’’ only. Scale Bar A-D = 50 μm Scale Bar E-E’’’ = 10 μm Scale Bar F-I’’ = 10 μm.

We first determined if the FA load of mitochondria is indeed reduced in the double mutants. Follicles incubated in media supplemented with C12:0-BODIPY displayed very small puncta in their NCs (Fig 5E-E’); these fluorescent puncta do not overlap with LD or mitochondrial staining (Fig 5E’’-E’’’). These observations show that in the double mutants, FAs no longer over-accumulate in mitochondria but are instead routed to a distinct compartment. A promising candidate is peroxisomes, as they have critical roles in some aspects of FA metabolism. By staining for the peroxisome marker PEX14 [37], we confirmed that in wild type, *CPT1* and *DGAT1* single mutants, and *DGAT1 CPT1* double mutants, peroxisomes are present in the NC cytoplasm, appearing as small puncta (Fig 5F-J). Next, we incubated explanted follicles with C12:0-BODIPY and then fixed and stained for peroxisomes. In wild type and the two single mutants, there was no detectable overlap between the two markers, but in the double mutant, PEX14 puncta and C12:0 puncta were superimposed (Fig 5H-H’’). These data suggest that in *DGAT1 CPT1* double mutants peroxisomes act as an alternative compartment to sequester FA and that FA accumulation in mitochondria is dramatically reduced.

This reversal of FA accumulation allowed us to interrogate the link between mitochondrial FA levels and mitoROS. The *DGAT1 CPT1* double mutants displayed reduced MitoSOX signal (S4A-D Fig), something we also observed when treating *DGAT1* follicles with etomoxir (S4E-G Fig). The double mutants also had lower MMP, probably due to the inability to use FAs for energy production; their MMP levels were indeed similar to those of the *CPT1* single mutant (Fig S5H-J). Because of the membrane potential dependence of MitoSOX, we used the ratio between MitoSOX and MMP to estimate mitoROS levels. The *DGAT1* mutants showed the highest ratio, while the double mutant, the *CPT1* single mutant, and wild type all showed similar ratios (S4K Fig). These data support the idea that the oxidative stress in *DGAT1* follicles is due to excessive amounts of mitochondrial FAs.

In the course of these studies, we noted that the mitochondria in our mutants exhibit morphological defects. In the wild type, NC mitochondria typically appeared as short rods (S4L Fig); in contrast, *DGAT1* mutants, *CPT1* mutants, and *DGAT1, CPT1* double mutants all displayed many toroidal NC mitochondria (S4L-P Fig). Toroidal mitochondria are often associated with mitochondrial stress, including oxidative stress [38–40]. Indeed, in wild-type follicles exposed to hydrogen peroxide, some mitochondria were also toroidal (S4N Fig). As not all of our mutants display increased mitoROS, these morphological changes cannot solely be due to oxidative damage but probably reflect additional insults to mitochondria, possibly linked to reduced MMP levels, which is shared between all these mutants.

### Throttling FA Influx into Follicles or Mitochondria Suppresses *DGAT1* Follicle Death

*DGAT1* mutant follicles die around S9 (Fig 6A-C) [9]. Is this follicle degeneration a consequence of the insult to mitochondria in this genotype, or due to some other aspect of dysregulated FA trafficking, or maybe just due to the absence of LDs? At S8, a nutrient checkpoint monitors the quality of follicles and can lead to developmental arrest [10, 41]. Lack of lipid-droplet accumulation might trigger this checkpoint, leading to follicle death, as previously suggested as an explanation for the apoptotic death observed for *DGAT1* mutants [9].

**Fig 6.**
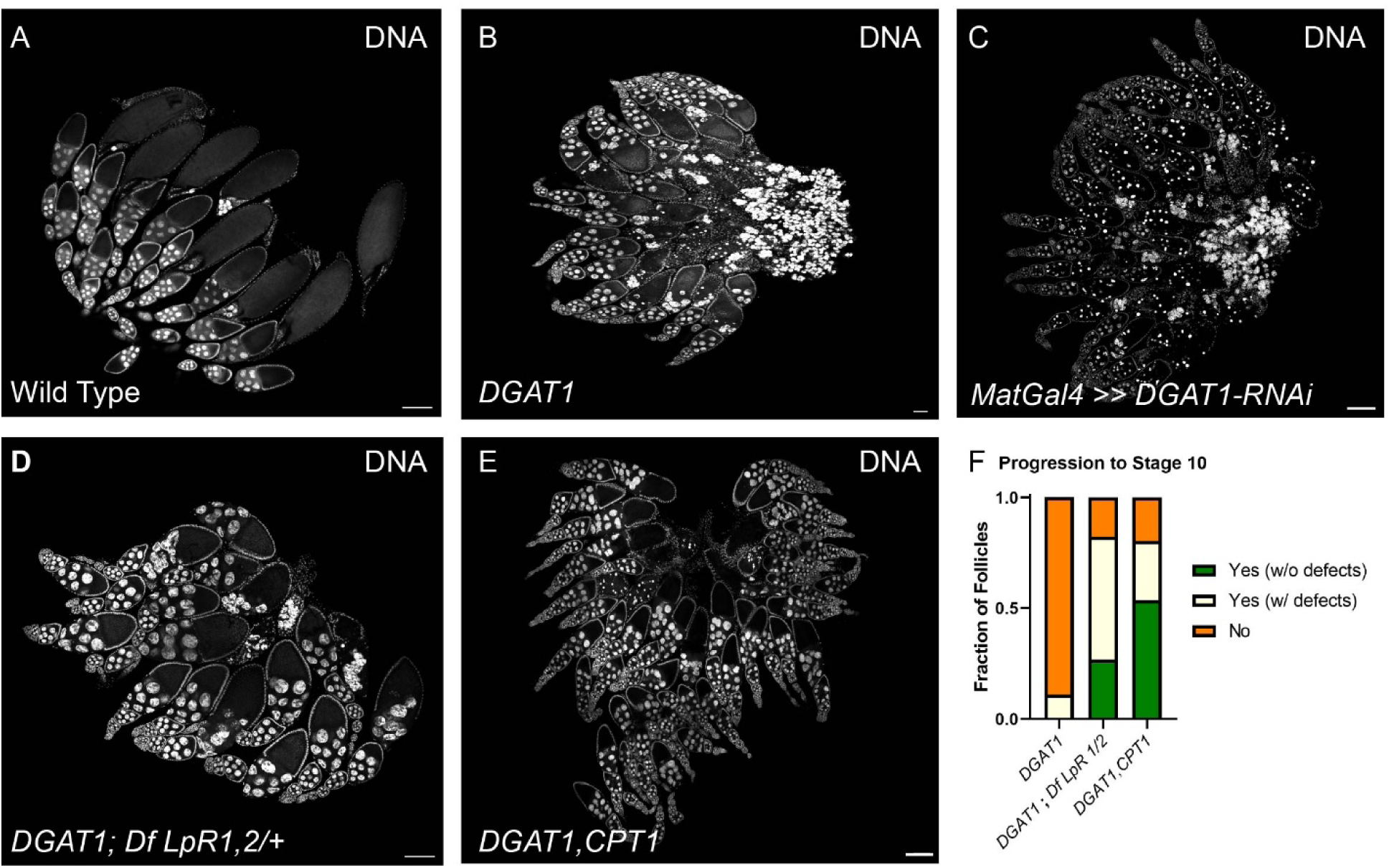
Throttling FA Influx into Follicles or Mitochondria Suppresses *DGAT1* Follicle Death. A-E) Tiled images of Hoechst-stained whole ovaries for wild type (A), *DGAT1* (B), *DGAT-RNAi* (C), *DGAT1*; *Df* LpR1,2/+ (D), and *DGAT1 CPT1* (E). F) Quantitation of follicle progression past stage 9 of oogenesis for the indicated genotypes. All images were acquired on Leica Stellaris (20x). Scale Bar A-E = 100 μm

As a first test, we throttled FA uptake into the follicle. In the wild type, FA uptake is mediated by lipophorin receptors LpR1 and LpR2. Using GFP-trap lines in the two genes, we found that, like in the wild type, LpR2 is expressed and present at the NC periphery in *DGAT1* mutants. LpR1 expression appears lower in wild-type follicles and is not readily detectable in the *DGAT1* mutants (S5A-B Fig). We then reduced the copy number of *LpR1* and *LpR2* in *DGAT1* mutants to decrease lipid uptake into the follicle. In this genotype, many follicles progressed past S9, with some reaching S10B (Fig 6D). To quantify this phenotype, we assigned follicles into three distinct categories: whether the oldest follicle in an ovariole was in S9 or had reached S10, either without or with defects. S10 follicles were scored to display defects if the oocyte length was shorter than the length of the NC cluster, if the follicular epithelium showed signs of impaired elongation, if NC nuclei displayed condensation, or if NCs exhibited membrane blebbing. Using this measure, the double mutant displayed a significant rescue (Fig 6F). Thus, FA influx into the follicle contributes to follicle death.

To determine if the death is specifically due to excess FA influx into mitochondria (as opposed to just influx into the follicle), we examined the *DGAT1 CPT1* double mutants. Numerous follicles progressed past S9, and some made it as far as S14 (Fig 6E, F). Quantitation using the criteria described above confirmed that these double mutants indeed progressed further into oogenesis (Fig 6F). These data support the idea that excessive FA influx into mitochondria causes developmental defects and triggers arrest.

It was previously shown that S9 *DGAT1* follicles undergo apoptosis as they show positive signal in TUNEL assays [9]. Similarly, we found that many S9 *DGAT1* mutants were positive for cleaved caspase 3, while wild-type S9 follicles were not (Fig 7A-B). *DGAT1* mutants with either reduced dosage of *LpR1* and *Lpr2* or lacking *CPT1* had relatively reduced incidence of cleaved caspase 3 signal in S9, consistent with their ability to develop further (Fig 7C-E).

**Fig 7.**
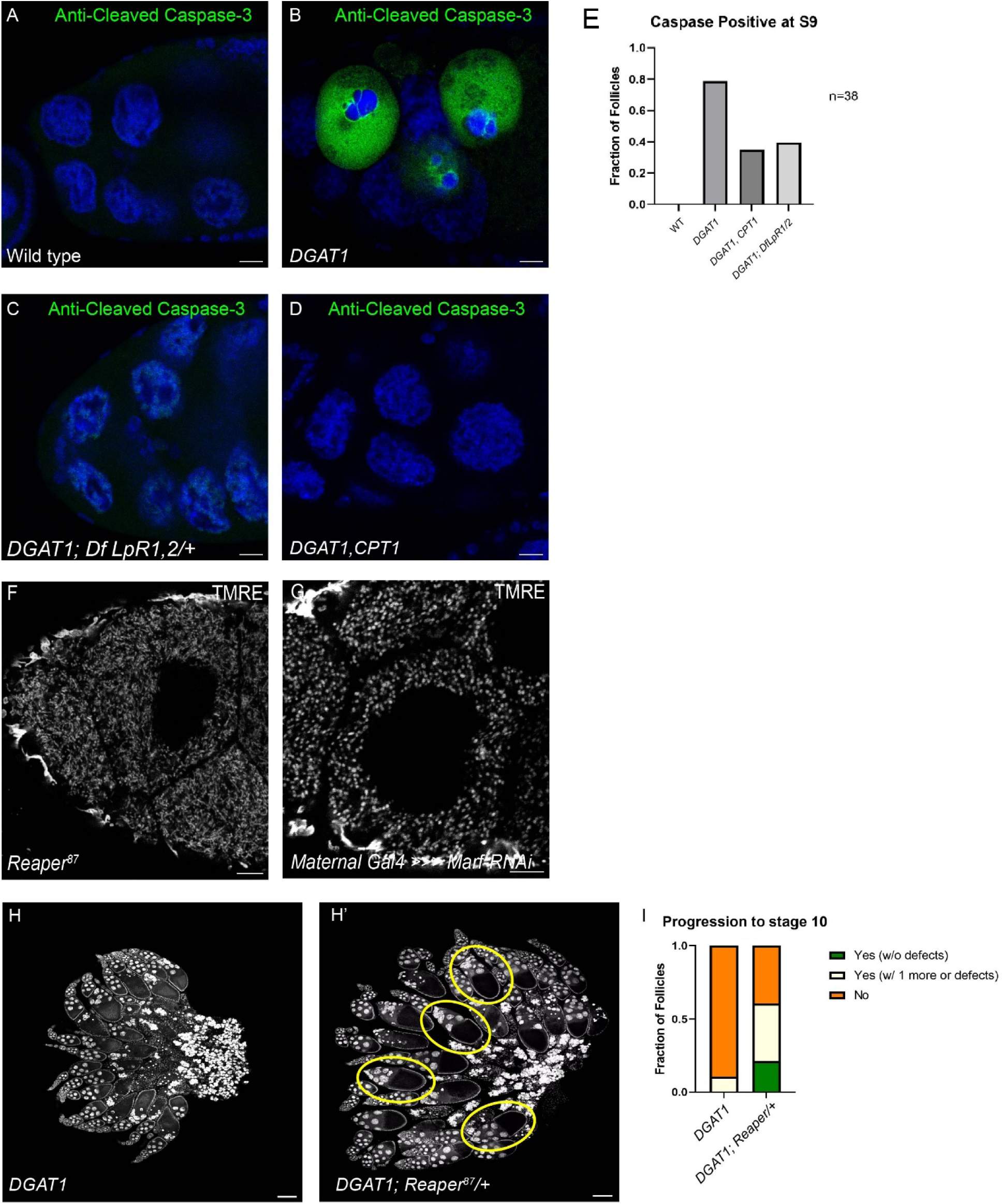
Damaged Mitochondria Contribute to Follicle Arrest in *DGAT1* mutants. A-D) Immunostaining with anti-cleaved caspase-3 antibody of wild-type (A), *DGAT1* (B), *DGAT1; Df* LpR1,2/+ (C), and D) *DGAT1, CPT1* (D) S9 follicles. E) Quantitation of S9 follicles that were positive for Caspase signal (n=38 for each genotype). F) Live image of a *Reaper^87^* S9 follicle after 15-minute incubation with TMRE (25nM). G) Live image of a *Marf-RNAi* follicle after 15-minute incubation with TMRE (25nM). The maternal tubulin Gal4 driver was used to drive the expression of *Marf-RNAi* in NCs. H-H’) Tiled images of Hoechst stained whole *DGAT1* (H) and *DGAT1*; *Reaper^87^*/+ (H’) ovaries. I) Quantitation of follicle progression past stage 9 of oogenesis for the indicated genotypes. Images were acquired on Leica Stellaris (20x) for panels H-H’ Scale Bar A-D = 10 μm. Scale Bar F-G 10 μm. Scale Bar H-H’ 100 μm.

Our data strongly suggest that in *DGAT1* mutant follicles damage to mitochondria leads to apoptosis and death. We therefore attempted to compromise the pathway for activating mitochondrial-induced apoptosis. The pro-apoptotic protein Reaper functions to inhibit mitochondrial fusion and blocks anti-apoptotic enzymes [42]. *Reaper^87^* null mutants have NC mitochondria that are elongated, likely due to increased activity of Marf, a mitochondrial fusion protein (Fig 7F-G) [42]. *DGAT1* mutant follicles with reduced *Reaper* dosage exhibited an increased frequency of follicles progressing past S9 into S10, suggesting that Reaper triggers mitochondrial-induced apoptosis in *DGAT1* mutants (Fig 7H-I). Overall, our data support the notion that NC mitochondria play a central role in the arrest phenotype of *DGAT1* mutants.

## Discussion

Our findings identify a new role for LDs in supporting *Drosophila* oogenesis. Previous studies had shown that LDs are important for regulating actin remodeling in S10B NCs and for achieving high levels of histones in oocytes. TAGs within NC LDs provide arachidonic acid to produce PGF_2ɑ_, which then activates a signaling pathway to drive the actin remodeling necessary for follicle morphogenesis [20]. In the oocyte, histones H2A, H2B, and H2Av are stored on LDs by the anchoring protein Jabba, which protects them from degradation [43]. In addition, Jabba-dependent histone sequestration to LDs occurs already in S9 and S10 NCs and likely promotes the transfer of histones to the oocyte [43]. Our data now reveal an essential metabolic role for NC LDs: during S9 and S10, they are crucial for funneling the correct amount of FAs to mitochondria for energy production. Without FA release from triglycerides in LDs, the MMP is reduced. If FAs cannot be stored in LDs, mitochondria are flooded with FAs, resulting in toxic levels of mitoROS that cause mitochondrial dysfunction, leading to follicle death. That the absence of LDs can lead to dysregulated FA trafficking and damage to mitochondria was previously described for cultured cells [26], but to our knowledge our study is the first instance to demonstrate this phenomenon in a whole tissue and to define its consequences for development.

### NC Mitochondria Rely on FAO to Produce Energy

Oogenesis is an energetically demanding process [1, 2], and in mammalian follicles, FAs have been identified as a key resource for energy production. However, there were no prior studies of whether FAs are utilized as a source of energy during *Drosophila* oogenesis. In fact, such a role appeared unlikely from mid-oogenesis onwards since it had previously been reported that S9 and 10 NC mitochondria are depolarized as they exhibited little to no TMRE fluorescence [15]. In contrast, we find significant TMRE incorporation at these stages (Fig 1); this incorporation is abolished when follicles are treated with the uncoupler FCCP to collapse the protein gradient across the mitochondrial membrane (Fig 1M, P), demonstrating that it indeed depends on the MMP. We suspect that the different results are due to the fact that the ex vivo TMRE incorporation was previously done in buffer [15], while we kept the follicles in nutrient-rich media to better approach physiological conditions.

Employing this experimental setup, we show that S9 and S10 follicles indeed engage in FAO and that it likely makes an important contribution to their energy metabolism. First, we find that the NCs exhibit FAOBlue signal, an assay that directly detects FAO activity (Fig 1A-A’’). This FAO signal is biologically meaningful as it is greatly diminished when we compromise FA import into mitochondria, either pharmacologically or genetically (Fig 1C-F). Second, the TMRE signal of these NCs is greatly diminished when import of FAs into mitochondria is impaired (Fig 1J-P), dropping by ∼50%. This drop is not an indirect effect of lost mitochondrial mass (S1C-D Fig) and thus represents reduced MMP. Finally, loss of CPT1 results in significant follicular defects, suggesting that mitochondrial FAO is crucial for follicle progression (Fig 1G-I).

### ATGL Activity Influences NC MMP

For LDs to provide FAs for FAO, the FAs must be released via lipophagy or via cytosolic lipases [44]. In many cells, the major cytosolic lipase is Adipose Triglyceride Lipase (ATGL) [45]. It was previously shown that ATGL is expressed in S10 NCs and necessary to release AA from LDs to support prostaglandin signaling [20]. The analysis presented here suggests that ATGL plays a much broader role, being also responsible for supplying mitochondria with FAs as fuel. Furthermore, loss of ATGL reduces both MMP and FAOBlue signal (Fig 3B-H). While our data cannot rule out that other lipases contribute to FA release in these follicles, they do suggest that ATGL plays a major role. In other cells, ATGL is heavily controlled by multiple different molecular mechanisms [46]. It will be interesting to determine in the future if any of these controls are active in follicles. We speculate that uncontrolled ATGL-mediated lipolysis could lead to lipotoxic overload of mitochondria, similar to what we observe in *DGAT1* mutants.

### How do FAs Exchange Between LDs and Mitochondria?

The data on ATGL strongly argue that FAs released from LDs are trafficked to mitochondria, but what paths these FAs take is not clear. Since FA transfer between organelles often occurs at membrane contact sites [19, 47, 48], we imagine two possible routes for such transfer in the follicle. First, the FAs might be directly transferred via LD-mitochondrial contacts [19]; second, there may be an indirect route via the ER, where FAs travel first through LD-ER contacts, then diffuse through the ER, and finally via ER-mitochondria contacts [49]. Unfortunately, neither of these contact sites are so far well characterized in *Drosophila*, and thus we cannot directly assess their distribution or functional involvement. However, in other systems, FA transfer via LD-mitochondria contacts is well established. For example, in mouse skeletal muscle, such LD-mitochondria contacts are regulated by the assembly of Rab8a-PLIN5 tethering complexes.

Here, Rab8a functions as a mitochondrial receptor for LDs by interacting with PLIN5, an LD surface protein. The tethering complex recruits ATGL to mobilize FAs from LDs to mitochondria [19]. Our imaging attempts have not revealed evidence for stable LD-mitochondrial contacts but also cannot rule them out: LDs and mitochondria undergo rapid streaming in the NC cytoplasm and move in and out of a given focal plane, so following their contacts over time is challenging. Future 3D live imaging of LDs and mitochondria may be required to resolve this issue. In any case, in S9 and S10 nurse cells, LDs, mitochondria, and ER are all highly abundant and broadly distributed throughout the NC cytoplasm. This arrangement implies that FAs have fairly short distances to travel between them, either through direct transient contacts between LDs and mitochondria or short ER stretches connecting the two organelles.

### Dysregulated Fatty Acid Trafficking in *DGAT1* Mutants Causes Mitochondrial Dysfunction

Since free FAs are known to induce lipotoxicity and cause tissue damage [26, 44, 50]; the trafficking of FAs must be tightly regulated. Thus, when cells are exposed to high levels of FAs, they are typically quickly esterified and sequestered into LDs. But how is damage to other cellular structures prevented before these FAs are safely incorporated into triglycerides? For NCs, our data suggest that protection may be achieved by minimizing the distance the FAs have to travel. Here, the incoming FAs come from lipophorin particles docked to LpRs at the nurse cell surface [13, 23]; from there, the FAs are presumably transported through the plasma membrane into the cell cortex. The cortex is rich in ER (Fig 2G-G’) and thus likely DGAT1: not only is DGAT1 known to reside in the ER [16], but we find that LDs are born at the cortex (Fig 2D-F). Thus, we propose that incoming FAs are turned into triglycerides almost immediately after they enter the NCs, thus minimizing the possibility that they reach other organelles and damage them.

Dysregulated FA trafficking by abolishing LD formation results in severe mitochondrial dysfunction. *DGAT1* mutant NC mitochondria are markedly depolarized, exhibit elevated ROS levels, and produce less ATP (Fig 4H-K). Elevated levels of mitochondrial FAs might have been expected to drive higher levels of FAO and thus increased MMP. However, our data revealed the opposite and argue that the drop in MMP is a secondary consequence of ROS damage, as incubation with the antioxidant NACA not only suppresses ROS levels but also increases MMP to near normal levels (S3A-D Fig, G-J). Mitochondrial dysfunction also occurs in cultured cells when DGAT1 function is abolished: starved mouse embryonic fibroblasts degrade membranes via autophagy and the released FAs, are then esterified into triglyceride by DGAT1 and sequestered within LDs; from here, ATGL mobilizes FA from LDs to fuel mitochondrial FAO. In the absence of DGAT1, free FAs are no longer sequestered into LDs, which dysregulates FA trafficking to mitochondria. This leads to an accumulation of toxic FAs and induces mitochondrial dysfunction. This metabolic routing of FAs to LDs serves as a protective buffer to control levels of free FAs [26]. Mitochondrial dysfunction can also occur under other conditions when FA influx is too high: in mouse skeletal muscle, a high-fat diet or insulin resistance can increase the expression of CPT1 and promote FA uptake into mitochondria, eventually exceeding their oxidative capacity [18] and leading to incomplete FAO and accumulation of toxic FAO intermediates.

Certain cell types can protect themselves against mitochondrial FA overload by exporting excess FAs. For example, hepatocytes can repackage and export excess FAs in the form of very low-density lipoproteins (VLDL) [18]. This option is likely not available to NCs as there is no evidence that they export FAs as part of their normal function; indeed, such export would diminish the nutrients delivered to the oocyte to support embryogenesis. We propose that NC instead use the strategy to sequester any excess FAs into LDs and to release only enough via regulated lipolysis to sustain sufficient FAO, thus preventing mitochondrial overload.

Mitochondria in *DGAT1* mutant follicles display increased mitochondrial ROS. Our data argue that this mitochondrial dysfunction is driven, at least in part, by excessive FA import. Mitochondrial ROS levels are alleviated when we throttle FA import by pharmacological or genetic inhibition of CPT1 (S4A-G Fig). Exactly how excess FAs cause ROS remains to be determined. In cultured cells*, DGAT1* inhibition leads to the accumulation of acylcarnitines, which are thought to be particularly toxic [26, 51]. Acylcarnitines are FA metabolites that allow acyl groups to be imported from the cytosol into mitochondria. CPT1 catalyzes the conversion of acyl-CoA and free carnitine to acylcarnitine. In our incubation assay, acylcarnitines would inherit the fluorophore from the FLFA, and thus the increased fluorescence we observe in mitochondria (Fig 4C) is consistent with acylcarnitine overaccumulation. Excess acylcarnitines disrupt the balance of free CoA, thereby depleting the pool of free CoA needed for the Krebs cycle, creating a metabolic bottleneck [18]. Acylcarnitines are also toxic in excess because they can accumulate within the mitochondrial matrix and inner membrane and induce uncoupling, which can result in mitoROS buildup [18]. Mitochondrial ROS production can then trigger oxidative damage to mitochondrial lipids, proteins, and DNA through lipid peroxidation cascades [52], exacerbating mitochondrial dysfunction.

### Fatty Acids Accumulate in Peroxisomes when LDs and Mitochondria are not Accessible

Our analysis of *DGAT1 CPT1* double mutants illustrates the metabolic flexibility of FA trafficking in *Drosophila* follicles. On the one hand, the dramatic rescue of oogenesis compared to the *DGAT1* single mutant clearly shows that it is the accumulation of excess FAs in mitochondria that induces developmental arrest and follicle death when DGAT1 is missing. On the other hand, rescue is puzzling because in the double mutant these toxic FAs are expected to flood the rest of the cell, potentially wreaking havoc on other cellular structures and triggering severe lipotoxicity. One might expect that such damage would severely compromise the follicles. Our data suggest that these potentially harmful FAs are directed to peroxisomes, which may either store them safely or detoxify them. We have observed a similar rerouting of FAs to peroxisomes in larval macrophages lacking DGAT1 (White et el., in revision). Peroxisomes are multifunctional organelles that participate in FA catabolism, cellular redox homeostasis, and function in close coordination with mitochondria [53]. Specifically, peroxisomes catabolize very long-chain fatty acids (VLCFAs), shortening them into FAs that can subsequently be utilized by mitochondria for energy production [53]. In *DGAT1 CPT1* mutants, peroxisomes readily sequester C12:0-BODIPY (Fig 5H). Peroxisomal accumulation was not observed in the other genotypes, likely because these FAs get shuttled to LDs and/or mitochondria instead. Initial attempts to generate *DGAT1 CPT1* mutants with a dosage reduction of essential peroxisome genes have not been successful because intermediate genotypes were sick or infertile. Thus, it remains to be determined to what extent this FA sequestration to peroxisomes indeed prevents widespread lipotoxicity.

### Health of NC Mitochondria Influences Developmental Competency

To prevent the production of low-quality oocytes, *Drosophila* ovaries employ several mechanisms to eliminate metabolically compromised follicles. In early oogenesis, a developmental checkpoint can be triggered by the detection of defects in germline cysts or nutrient shortage, which leads to cell death in region 2 of the germarium [10]. A mid-oogenesis checkpoint monitors the overall health of the follicle and nutrient availability [10] and induces cell death in S8 to eliminate the follicles; it is thought to prevent wasting energy on poor-quality follicles before the energy-intensive final growth stages. There is also a developmentally controlled cell death pathway in S13 that gets rid of the NCs once they have transferred their contents to the oocyte [10]. Intriguingly, the death of *DGAT1* mutant occurs during S9 and thus does not correspond to any of the known cell death checkpoints, nor the developmentally induced apoptosis.

Our data indicate that follicle death in *DGAT1* mutants is due in large part to a mitochondrial quality control pathway. In the absence of DGAT1, NC mitochondria exhibit FA overload, partial depolarization, and elevated ROS production. When these insults to mitochondria are relieved through CPT1 inhibition, follicle death is dramatically delayed (Fig 6). In addition, reducing the pro-apoptotic protein Reaper mitigates the arrest phenotype (Fig 7); apoptosis signaling via Reaper usually involves its translocation to mitochondria, where it inhibits the anti-apoptotic protein DIAP1 and thus leads to caspase activation. Thus, the involvement of Reaper in *DGAT1* follicle death reinforces the notion that this death involves mitochondria.

Why would it be necessary for S9 follicles to assess the health of mitochondria? During S9 and S10, follicles undergo massive growth, show abundant levels of transcription and translation, and rearrange their cytoskeleton to prepare for the transfer of nurse cell contents to the oocyte. All these processes require large amounts of energy. Thus, if these follicles experience mitochondrial dysfunction and energy shortage, an eventual oocyte – if produced at all – would likely be of poor quality. It may therefore be more beneficial to destroy a follicle thus compromised rather than invest more resources into it and generate a defective oocyte.

Alternatively, or in addition, this quality control mechanism may safeguard against transferring cytotoxic materials to the oocyte.

## Materials and Methods

### Fly Stocks

All stocks used in the experiments were maintained on a standard brown fly food medium. Food was prepared by combining the following ingredients per liter of deionized water: agar (7.9 g/L), molasses (22 g/L), malt extract (75 g/L), brewer’s yeast — brown (18 g/L), corn flour (80 g/L), soy flour (10 g/L), methyl paraben (2 g/L), ethanol (7.2 ml/L), and propionic acid (6.3 ml/L). Flies for experiments were kept at room temperature unless noted. The following stocks were used: *Oregon R* (BDSC #5), *DGAT1^QX25^ (midway)* (BDSC #5095), *CPT1^1^* (*whd*) (BDSC #441), *LpR1-GFP* (BDSC #60145), *LpR2-GFP* (BDSC #60219), *ATGL^1^* (bmm) ([34]), ATGL-RNAi (BDSC # 25926), DGAT1-RNAi (BDSC #65963), Lpr1-,LpR2-/Tm3 were generated by Joaquim Culi’s group [23], Reaper^87^ (BDSC #83150), PDI-GFP (BDSC #6839), UAS-Marf-RNAi (BDSC #67158). The *DGAT1 CPT1* double mutant chromosome was generated by recombination. UASz-ATPSnFR flies were generated from the ‘cyto-Ruby3-iATPSnFR1.0’ vector described in Lobas *et al.* 2019 [36]. The construct was transferred into the UASz1.1 [54] vector using standard restriction enzyme cloning with Xho1 and BamH1. This vector was then inserted at the attP2 site on chromosome 3 by BestGene.

### CPT1 Outcross Strategy

The *CPT1^1^* allele is located on the second chromosome and contains a 16bp deletion in exon 2; this deletion results in a frameshift and a truncated protein. Adults homozygous for *CPT1^1^* were previously described as having crinkled wings [25]. We confirmed by genomic PCR and sequencing that flies from BDSC stock #441 do carry the deletion; however, their wings are indistinguishable from wild type. After outcrossing and re-isolating multiple second chromosomes, the wing phenotype returned in some progeny, suggesting that the original stock had accumulated suppressors. We picked one stock that had both the 16 bp deletion and the wing phenotype and used it as the *CPT1* mutant for our analysis.

### Dissection Protocol for Fixed Tissue

Adult female and male flies (to allow for mating), younger than 2 weeks old, were fed dry yeast for 48 hours at room temperature to promote mating and egg production. Ovaries were dissected in Schneider’s *Drosophila* medium (Sigma-Aldrich) and fixed for 12 minutes at room temperature in 4% paraformaldehyde diluted in 1× phosphate-buffered saline (PBS). Following fixation, samples were washed three times for 10 minutes each in PBS containing 0.1% Triton X-100 (PBT). After their respective staining period, follicles were counterstained with Hoechst 33258 (1µg/mL) for 20 minutes and washed with PBT prior to mounting.

### Dissection Protocol for Live Tissue

Adult female and male flies (to allow for mating), younger than 2 weeks old, were fed dry yeast for 48 hours at room temperature to promote mating and egg production. Ovaries were dissected in maturation medium (Schneider’s *Drosophila* medium, Sigma-Aldrich; 15% FBS, Gibco; 10 mg/ml insulin, Sigma Aldrich; 1× penicillin/streptomycin (10,000 U/ml penicillin and 10 mg/ml streptomycin, pH 6.95), Gibco), and individual follicles were separated from the ovary/muscle sheath and transferred to an Eppendorf tube for treatment with a reagent. This

### Live Imaging

Following dissection and any indicated pharmacological treatments, individual follicles were transferred from the Eppendorf tube to a poly-L-lysine-coated glass-bottom MatTek imaging dish containing 200 µL of maturation medium, ensuring that each follicle was fully submerged throughout imaging. Live imaging was performed immediately after transfer using either a Leica SP5 confocal microscope or a Leica STELLARIS confocal microscope equipped with an HC PL APO 40×/1.30 Oil CS2 objective. Imaging was conducted on a temperature-controlled stage maintained at 25°C under ambient atmospheric CO₂ conditions. Identical image acquisition settings were maintained across experimental groups within each experiment, and follicles remained immersed in maturation medium for the duration of image acquisition.

### Immunofluorescence

Following the dissection protocol for fixed tissue, ovaries were blocked overnight at 4°C in a blocking buffer consisting of 10% bovine serum albumin (BSA), 0.1% Triton X-100, and 0.02% sodium azide in PBS (Ovary block). Ovaries were washed with PBT (1x PBS, 0.1% Triton X-100) three times for 10 minutes and then blocked in ovary block (10% bovine serum albumin, 0.1% Triton X-100, 0.02% sodium azide in PBS) overnight at 4°C. Primary antibody incubation was performed using rabbit anti–Cleaved Caspase-3 (Asp175; Cell Signaling Technology, cat. no. 9661) diluted 1:400 in ovary block. After three washes in PBT, samples were incubated overnight at 4°C with goat anti-rabbit IgG conjugated to Alexa Fluor 488 (1:750 dilution in blocking buffer; Thermo Fisher Scientific). Ovaries were washed three additional times in PBT and mounted in Aqua-Poly/Mount (Polysciences) for imaging.

### Lipid Droplet Staining for Fixed Tissue

Following the dissection protocol for fixed tissue, ovaries were transferred to an Eppendorf tube with PBT and were mechanically dissociated using a pipette until follicles separated from each other. For Nile red staining, a 1 mg/ml stock was prepared using acetone and added to fixed ovaries (1:50) in ovary block to incubate for 1 h at room temperature. For LipidSpot staining, follicles were incubated in Ovary block with LipidSpot (1:100) for 20 minutes at room temperature. Afterwards, follicles were washed three times for 10 minutes each in PBT. Samples were then mounted in Aqua-Poly/Mount and imaged.

### Ex-Vivo Fluorescent lipid species Incubation

Following **Dissection Protocol for Live Tissue**, follicles were isolated from the ovary and incubated for 15 minutes in maturation medium supplemented with 5 µM fluorescent lipid species (S1 Table 1). The medium was removed by aspiration and replaced with fresh maturation media. The follicles were then transferred to a poly-D-lysine-coated, glass-bottom culture dish with 200 µl of maturation medium. Follicles were allowed to adhere to the culture dish for 5 min prior to imaging.

### Feeding FLFA-Supplemented Yeast Paste

Flies were fed supplemented yeast paste (1% sucrose, 5 µM fluorescent fatty acid, dry yeast) overnight, and follicles were dissected, fixed, and stained for LDs using LipidSpot 610 (1:100, Biotium). In some cases, LipidSpot 610 was omitted and LDs were detected by confocal reflection microscopy [30, 55] instead.

### Image Acquisition

Microscope images of fixed and live *Drosophila* follicles were obtained using the following microscopes: Leica SP5 confocal microscope and Leica Stellaris. For the Leica SP5 confocal, an HCX PL APO CS 40x/1.40 oil UV objective was used for image acquisition. For the Leica Stellaris, an HC PL APO CS2 63x/ 1.40oil objective or HC PL APO 20×/0.75 IMM CORR CS was used for image acquisition. The images were acquired using the Leica SP5 unless otherwise stated. Post-acquisition analysis was performed using Fiji/ImageJ. Quantitation was conducted using FIJI’s integrated density measurement tool and ROI manager. Adobe Illustrator was used to assemble figures. GraphPad Prism was used to determine P values and make graphs.

### Quantitation of Oogenesis Defects

We manually staged follicles based on the criteria in [8]. We scored the following signs of developmental arrest: 1) an abnormal ratio between oocyte length and NC size, such that the oocyte length is not the same or close to the same in length as the NCs (i.e., the ratio is no longer 1:1); 2) impaired elongation of the follicular epithelium; 3) condensation of NC nuclei, and 4) membrane blebbing of NCs. For each ovariole, we examined the most mature follicle and assigned it into one of three categories: “no” if: follicles arrest at S9 and show multiple signs of arrest; “yes with defects” if follicle had progressed past S9 but exhibited sign(s) of arrest; “yes” if follicle progressed past S9 with no obvious sign of arrest. The analysis of follicular arrest was done using Fiji/ImageJ line tool to measure the dimensions of follicles, and NC nuclei size was measured to decide whether nuclei had condensed.

**Fig 8.**
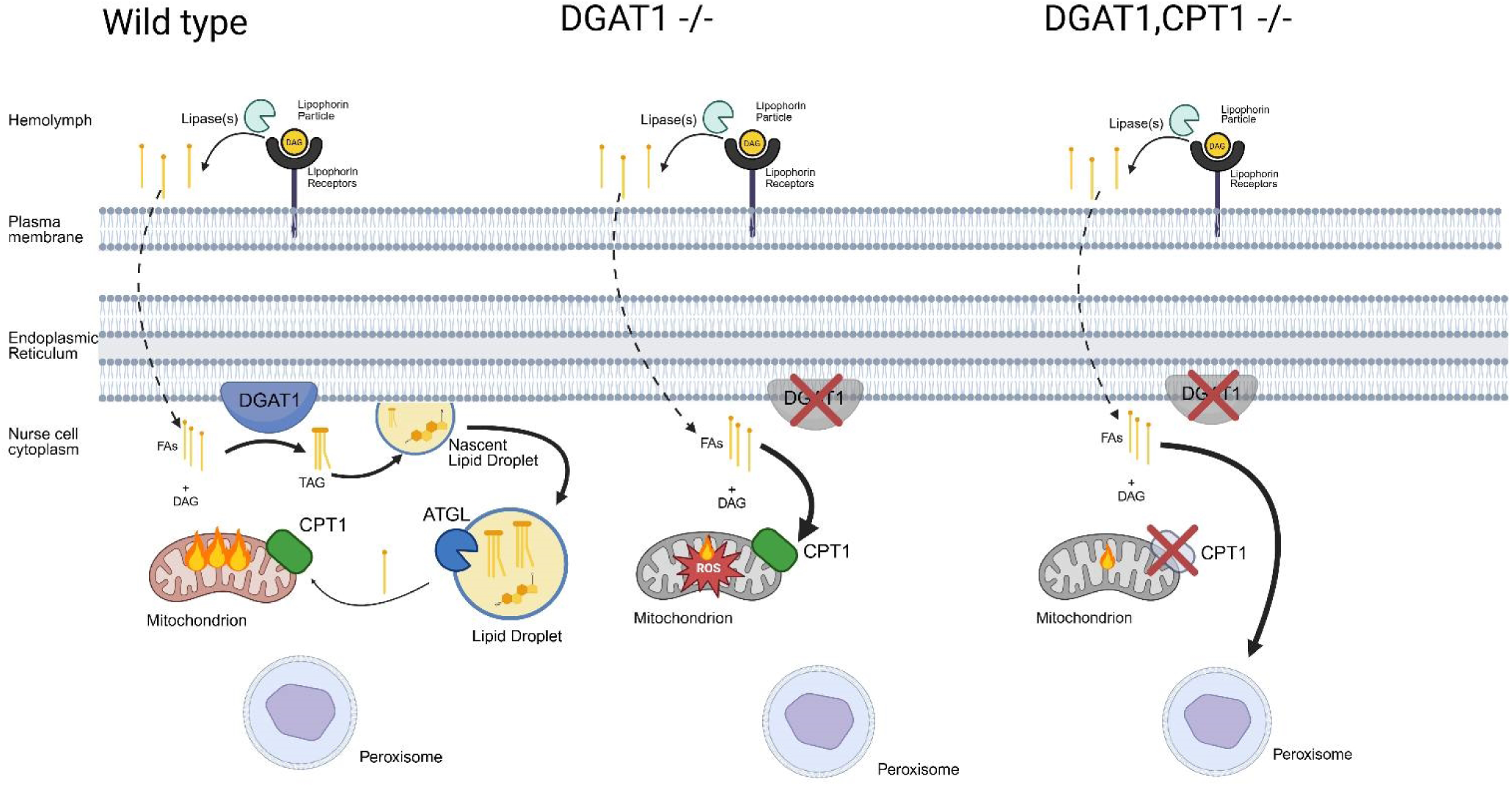
Model of Fatty Acid Trafficking during Mid-Oogenesis and the Role of LDs in Protecting Mitochondrial Integrity. A) Wild type. Beginning at stage 9 of oogenesis, lipid is rapidly taken up from the hemolymph in the form of DAG-rich lipophorin (Lpp) particles. Lpp particles dock onto lipophorin receptors at the plasma membrane, where their contents are hydrolyzed by lipases and free fatty acids (FAs) are taken up by nurse cells (NCs). The endoplasmic reticulum (ER) lines the periphery of the NC, where incoming FAs are quickly esterified by DGAT1 to form triacylglycerol (TAG), which accumulates between the ER leaflets until a nascent lipid droplet (LD) buds off. Mature LDs serve as a dynamic FA reservoir. ATGL releases FAs from LDs, and the subsequent uptake of long-chain FAs into mitochondria is mediated by CPT1. Once imported, FA undergoes β-oxidation to generate ATP. Tight regulation of FA trafficking from the hemolymph through nurse cells to mitochondria is essential for the completion of oogenesis and the production of competent follicles. B) *DGAT1* mutant follicles. In the absence of DGAT1, FA is taken up by the follicle normally but cannot be stored in LDs. As a result, FA is excessively redirected to mitochondria, leading to mitochondrial FA overload and accumulation of reactive oxygen species (ROS). ROS negatively affects mitochondrial membrane potential and ATP synthesis, contributing to the initiation of apoptosis and eventual follicular degeneration. C) *DGAT1 CPT1* follicles. When both DGAT1 and CPT1 are non-functional, FAs that can no longer be stored in LDs or imported into mitochondria are instead shunted to peroxisomes. Reducing mitochondrial FA overload in this way partially rescues the follicular arrest phenotype; however, double mutants still fail to produce fertilization competent follicles.

## Acknowledgements

We thank Andrew Simmonds from the University of Alberta for sharing the anti-Pex14 antibody. Stocks obtained from the Bloomington *Drosophila* Stock Center (NIH P40OD018537) were used in this study. We are grateful to Tina Tootle for feedback and advice on various aspects of this study. Figure 8 was generated with BioRender (Created in BioRender. Whi, R. (2026) https://BioRender.com/km6tj55). This work was supported by NIH grants R01 GM102155 to MAW and 1F31 HD100127 to MDK.

## Supporting Information

**S1 Fig.**
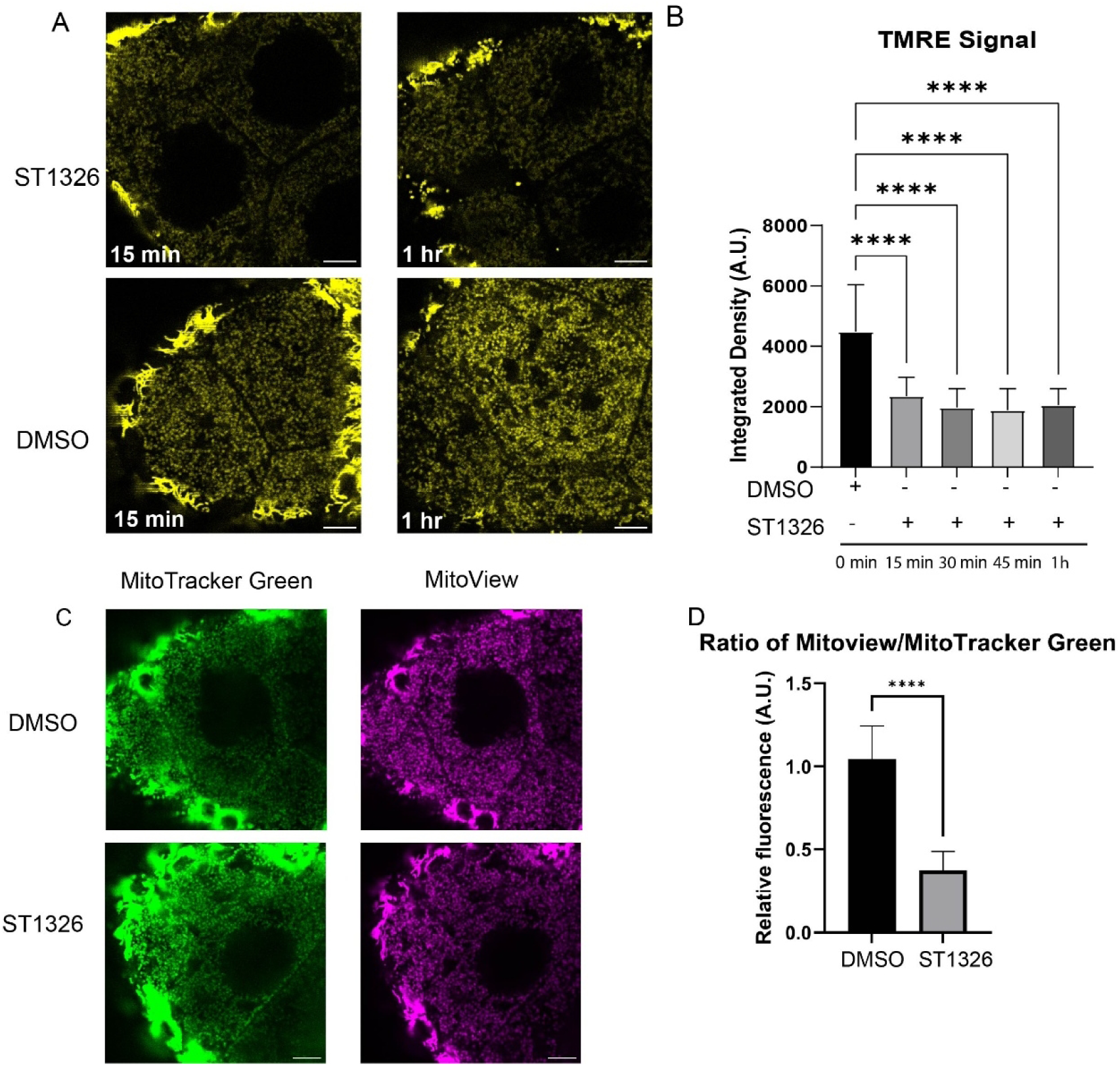
Pharmacological Inhibition of CPT1 Reduces Membrane Potential. A) Live images of wild-type S9 follicles treated with ST1326 or DMSO vehicle control, captured at 15 minutes and 1 hour post-treatment. Follicles were stained with TMRE (25 nM) to assess mitochondrial membrane potential. B) Quantitation of TMRE integrated density over time in ST1326- and DMSO-treated S9 follicles. ST1326 treatment causes a significant reduction in TMRE signal relative to DMSO controls at all timepoints examined. C) Wild-type S9 follicles stained with MitoTracker Green (mitochondrial mass marker) and MitoView (mitochondrial membrane potential dye) following DMSO or ST1326 treatment. D) Quantitation of MitoView signal normalized to MitoTracker Green signal in S9 nurse cells, showing a significant reduction in membrane potential per unit mitochondrial mass in ST1326-treated follicles relative to DMSO controls. Data represents SD. ****p < 0.0001. Scale bars: 10 μm (A, C).

**S2 Fig.**
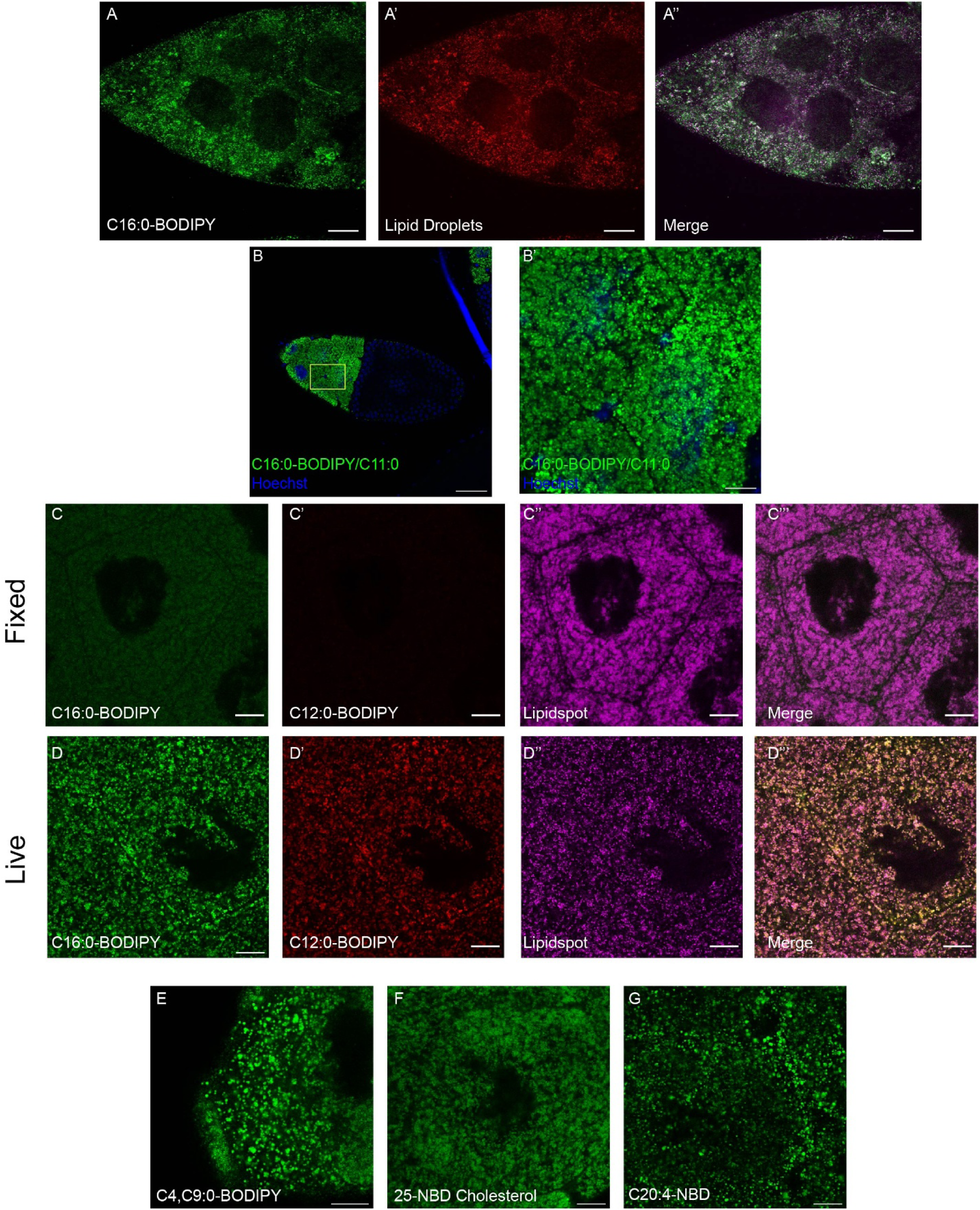
Visualizing Sequestration of Fluorescent Fatty Acid by NC LDs. A-A’’) Live images of wild-type S9 follicle from adult female that had been feed C16:0-BODIPY yeast paste for 20 hours. BODIPY signal (A), LDs detected by confocal reflection microscopy (CRM) (A’), merge of the two channels. B) Wild-type follicle supplemented with C16:0-BODIPY/C11:0 (DAG) supplemented WT follicle and stained with Hoechst. B’) Magnified region of the follicle from B. C-D’’) Wild-type S9 nurse cells exposed to C16:0-BODIPY (C, D) and C12:0-BODIPY (C’, D’) and stained with LipidSpot (C’’, D’’). C’’’ and D’’’ show merge of all channels. The follicle in (C-C’’’) was fixed prior to supplementation with fluorescent FAs. The follicle in D-D’’’ was alive during incubation with fluorescent FAs. E) Wild-type nurse cell after incubation with C4:0-BODIPY, C9:0-BODIPY. F) Wild-type nurse cell after incubation with 25-NBD cholesterol. G) Wild-type nurse cell after incubation with C20:4-NBD. Scale Bar A-A’’ = 20μm. Scale Bar B and B’ = 50μm. and 10μm respectively Scale Bar C-G = 10μm.

**S3 Fig.**
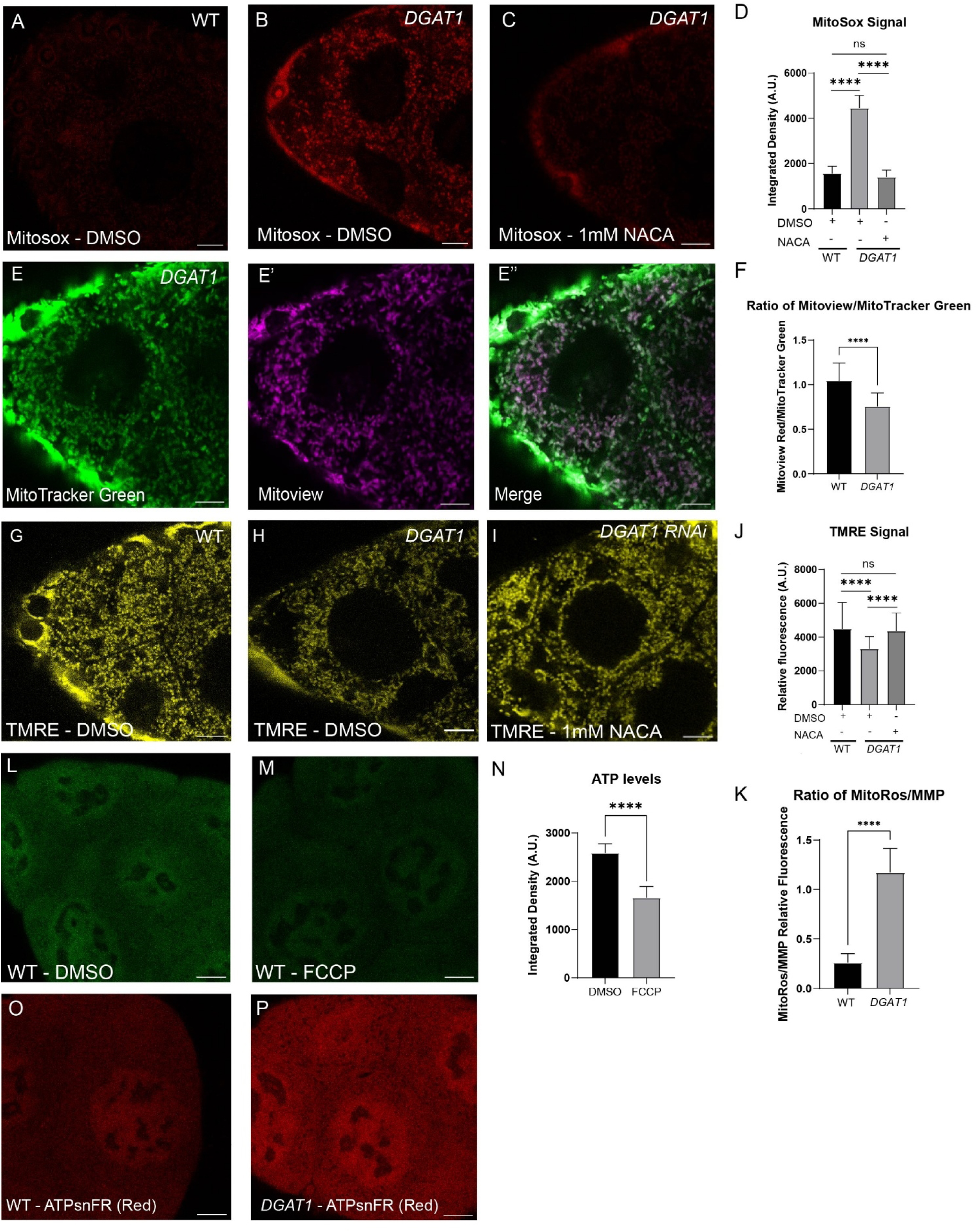
*DGAT1* Mutants Display Multiple Mitochondrial Defects. **A)** Live image of MitoSox staining for wild-type, B) *DGAT1*, and C) NACA-treated *DGAT1* mutant S9 follicles. D) Quantitation of MitoSox signal for the indicated conditions. E-E’’) Live images of the same *DGAT1* follicle stained with MitoTracker Green (E) and MitoView (E’). E’’ shows merge of both channels. F) Quantitation of the ratio of MitoView and MitoTracker Green relative fluorescence for the indicated genotypes. G-I) Live images of TMRE (25nM) staining for wild-type (G), *DGAT1* (H), and NACA-treated *DGAT1* RNAi (I) S9 follicles. J) Quantitation of TMRE signa for the indicated conditions. K) Quantitation of the ratio of MitoRos to mitochondrial membrane potential (MitoRos/MMP) for wild-type and *DGAT1* mutant S9 follicles. L, M) Live images of ATP synthase sniffer (ATPsnFR) signal in nurse cells of wild-type S9 follicles, untreated (L) or treated with FCCP. N) Quantitation of ATP synthase sniffer signal. O, P) Red channel of ATPsnFR in wild-type and *DGAT1* nurse cells used to normalize green channel signal. Scale Bar A–C, E–E’’, G–I, L–M, O–P = 10μm.

**S4 Fig.**
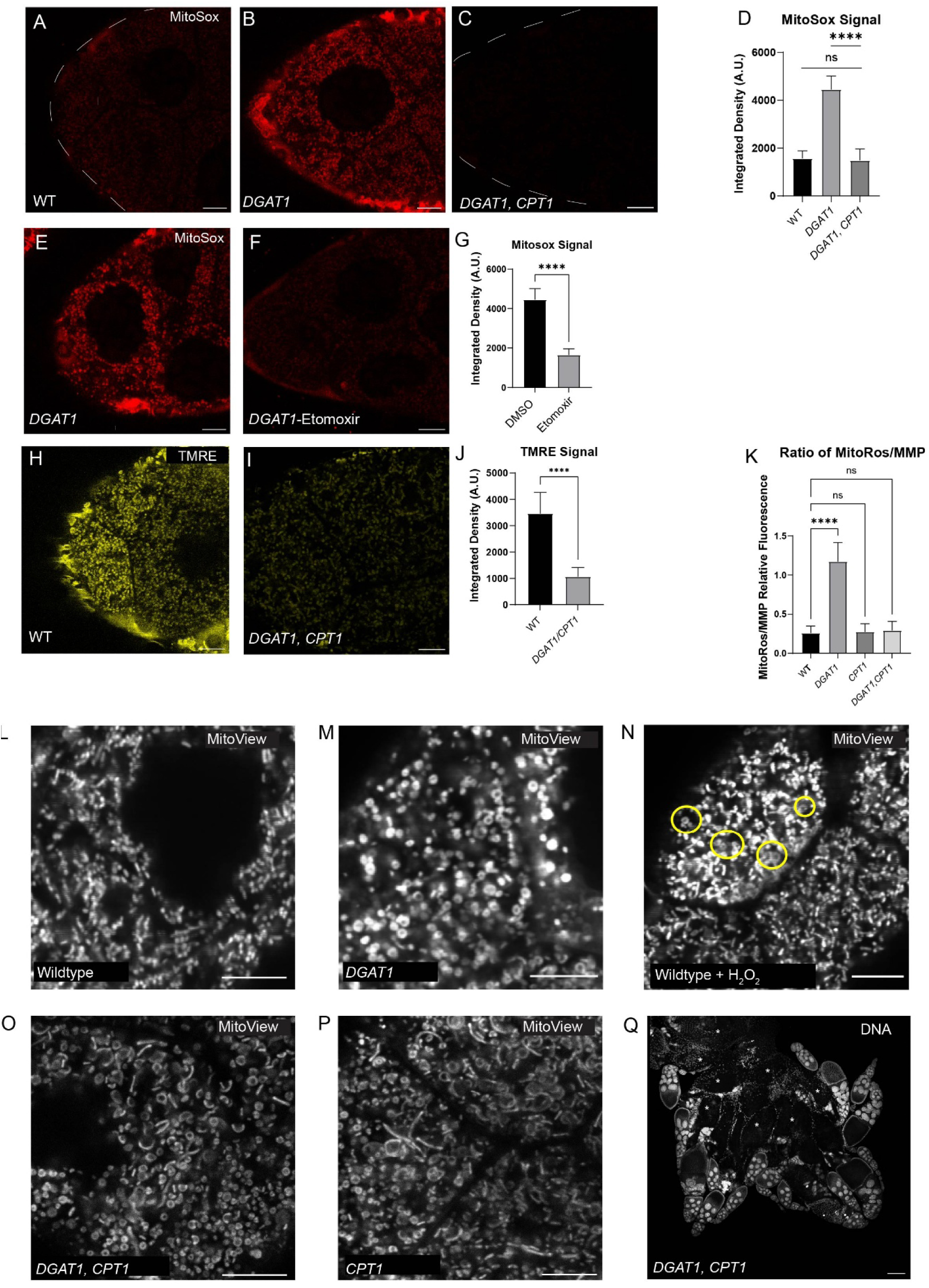
Perturbations to DGAT and CPT1 Function Affect both Mitochondrial Bioenergetics and Morphology. A–C) Representative confocal images of MitoSox staining in wild-type (A), *DGAT1* mutant (B), and *DGAT1 CPT1* double mutant S9 follicles (C). D) Quantitation of MitoSox integrated density in NCs of S9 follicles across genotypes. *DGAT1* mutants show a significant elevation in MitoSox signal relative to wild type, while signal in *DGAT1 CPT1* double mutants is not significantly different from *DGAT1* single mutants. E–F) Representative confocal images of MitoSox staining in *DGAT1* mutant S9 follicles treated with DMSO vehicle (E) or Etomoxir (F). G) Quantitation of MitoSox integrated density, demonstrating significant reduction of ROS signal following Etomoxir treatment in *DGAT1* mutants. H–I) Representative confocal images of TMRE staining in wild-type (H) and *DGAT1, CPT1* double mutant (I) follicles. J) Quantitation of TMRE integrated density in S9 follicles, demonstrating significantly reduced mitochondrial membrane potential in *DGAT1 CPT1* double mutants relative to wild type. K) Quantitation of the ratio of MitoROS signal to mitochondrial membrane potential (MitoROS/MMP) in S9 follicles across genotypes. L–P) Representative confocal images of MitoView staining in nurse cells from wild-type (L), *DGAT1* mutant (M), wild-type treated with H₂O₂ (yellow circles indicate fragmented mitochondria) (N), *DGAT1 CPT1* double mutant (O), and *CPT1* mutant (P) S9 follicles. Q) Hoechst staining of a *DGAT1 CPT1* whole ovary. Data represent mean ± SD. ****p < 0.0001; ns, not significant. Images were acquired on Leica Stellaris (20x) for panel Q only. Scale bars: 10 μm (A–C, E–F, H–I, L–P).

**S5 Fig.**
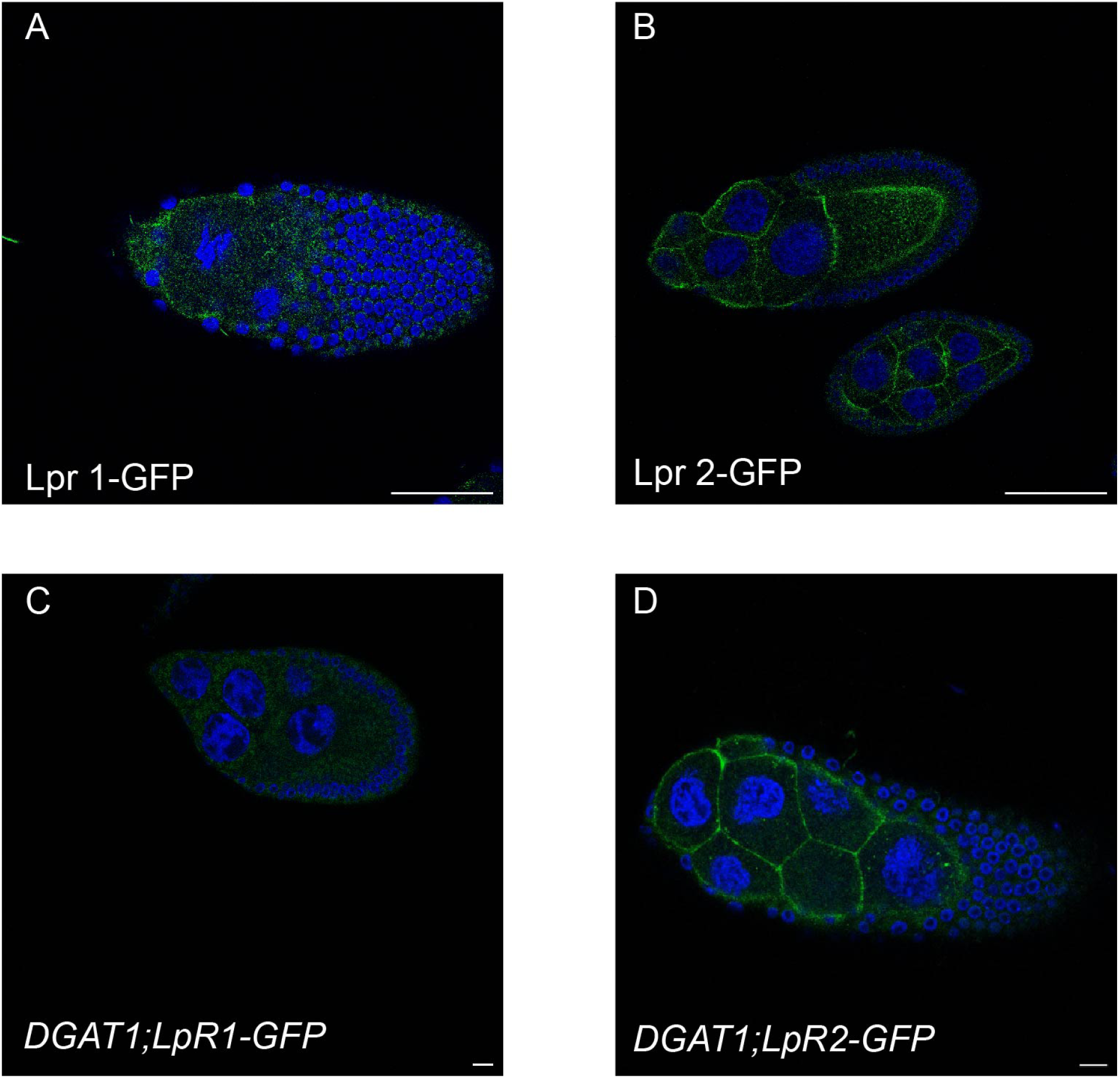
Visualizing LpR Expression during Mid-oogenesis. LpR1- or Lpr2-GFP expressing S9 follicles stained with Hoechst (blue) to detect nuclei. **A**) LpR1-GFP S9 follicle. B) LpR2-GFP S9 follicle. C) *DGAT1*; LpR1-GFP S9 follicle. D) *DGAT1*;LpR2-GFP S9 follicle. Scale Bar A-B = 50μm C-D = 10μm.

## Supplemental Movies

S1 Video. Real-Time Visualization of C16:0-BODIPY incubation. **A)** Image sequence of wild-type S10 follicle in maturation media supplemented with 5 µM C16:0-BODIPY (green). **B)** Image sequence of wild-type follicle in maturation media supplemented with 5 µM C16:0-BODIPY (green). CRM was utilized to visualize LD (magenta). Frame rate: 250 frames, 17.86 sec. Total duration: 58 minutes. Scale bar = 50 µm

S2 Video. Real-Time Visualization of C12:0-BODIPY incubation in PDI-GFP S10 follicles. **A)** Image sequence of PDI-GFP nurse cells in maturation media supplemented with 5 µM C12:0-BODIPY. **B)** CRM was used to visualize LDs. **C)** PDI-GFP expression. **D)** Merge of C12:0-BODIPY and CRM. The follicle was not treated with any drug prior to imaging. Frame rate: 16 frames, 4 sec. Total duration: 40 minutes. scale bar 10 µm

S3 Video. Colchicine Treatment of C12:0-BODIPY-Supplemented PDI-GFP S10 Follicle. **A)** Prior to imaging, the follicle was treated with 80µg/mL colchicine for 1 hour in maturation media. PDI-GFP follicle was then supplemented with 5 µM C12:0-BODIPY in maturation media. **B)** CRM was used to visualize LDs. **C)** PDI-GFP expression. **D)** Merge of C12:0-BODIPY and CRM. Frame rate: 15 frames, 3.75 sec. Total duration: 37.5 minutes. Scale bar 10 µm

**S1 Table.** Fluorescent lipid species.

| Lipid Species | Company | Saturation |
| --- | --- | --- |
| BODIPY™ 500/510 C4, C9 (5-Butyl-4,4-Difluoro-4-Bora-3a,4a-Diaza-s-Indacene-3-Nonanoic Acid)) | Thermo Fisher Scientific | Saturated |
| BODIPY™ FL C12 (4,4-Difluoro-5,7-Dimethyl-4-Bora-3a,4a-Diaza-s-Indacene-3-Dodecanoic Acid)<br>BODIPY™ FL C16 (4,4-Difluoro-5,7-Dimethyl-4-Bora-3a,4a-Diaza-s-Indacene-3-Hexadecanoic Acid) | Thermo Fisher Scientific | Saturated |
| 1-palmitoyl-2-(dipyrrometheneboron difluoride) undecanoyl-sn-glycerol | Avanti Polar Lipids | Saturated |
| 25-[N-[(7-nitro-2-1,3-benzoxadiazol-4-yl)methyl]amino]-27-norcholesterol | Avanti Polar Lipids | N/A |
| 20-[(7-nitro-2-1,3-benzoxadiazol-4-yl)amino]arachidonic acid | Avanti Polar Lipids | Polyunsaturated |

